# Human-like telomeres in *Zostera marina* reveal a mode of transition from the plant to the human telomeric sequences

**DOI:** 10.1101/2020.03.11.987156

**Authors:** Vratislav Peska, Martin Mátl, Terezie Mandákova, Daniel Vitales, Petr Fajkus, Jiří Fajkus, Sònia Garcia

## Abstract

A previous study describing the genome of *Zostera marina*, the most widespread seagrass in the Northern hemisphere, revealed some genomic signatures of adaptation to the aquatic environment. Important features related to the ‘back-to-the-sea’ reverse evolutionary pathway were found, such as the loss of stomatal genes, while other functions like an algal-like cell wall composition were acquired. Beyond these, the genome structure and organization were comparable to the majority of plant genomes sequenced, except for one striking feature that went unnoticed at that time: the presence of human-like instead of the expected plant-type telomeric sequences. By using different experimental approaches including FISH, NGS and Ba131 analysis, we have confirmed its telomeric location in the chromosomes of *Z. marina*. We have also identified its telomerase RNA subunit (TR), confirming the presence of the human-type telomeric sequence in the template region. Remarkably, this region was found to be very variable even in clades with a highly conserved telomeric sequence across their species. Based on this observation, we propose that alternative annealing preferences in the template borders can explain the transition between the plant and human telomeric sequences. The further identification of paralogues of TR in several plant genomes brought us to the hypothesis that plants may keep an increased ability to change their telomeric sequence. We discuss the implications of this occurrence in the evolution of telomeres while introducing a mechanistic model for the transition from the plant to the human telomeric sequences.

---

*Zostera marina*, or common eelgrass, belongs to the family Zosteraceae, one of the four Alismatales families (basal monocots) that make up the seagrasses. *Zostera marina* plays a crucial role in coastal ecosystems around the world, providing food and shelter to numerous species. Although it is considered of least concern in terms of conservation status, the population trend is decreasing worldwide (IUCN red list), threatened by processes such as fishing, pollution, and the presence of invasive species^1,2^. Recently, the genome of this species was reported^3^, a resource that has already aided in functional ecological studies in seagrass ecosystems under global warming^4,5^ or evolutionary research^6,7^.

## Plants with unusual telomeres

A fundamental trait of a species genome is the composition of its telomeres. Telomeres are terminal chromosomal domains, whose minisatellite sequence is well conserved across large groups of organisms. Basically, we can refer to the plant-type telomere repeat as (TTTAGGG)n, shared by most plants, and to the human-type telomere repeat as (TTAGGG)n, shared by vertebrates and some other animals, while more diverse repeats can be found in fungi, insects, and others. Such typical telomeric sequences are mostly taken for granted in any species from a given group, yet recent experience prevents us from making such generalizations. For the discovery of unusual telomeric sequences in *Cestrum* and *Allium* (well-known genera for their ornamental and crop plants, respectively), which had remained enigmatic for decades, the availability of NGS data and bioinformatics tools have been crucial, together with molecular cytogenetics^8^. The finding of unusual telomeric sequences have had a great scientific impact, for instance, the exceptionally long *Allium* sequence motif^9^ was further utilized as bait for the identification of *bona fide* telomerase RNAs (TRs) in plants^10^.

## *Zostera marina* has human-type telomeres

Here we provide compelling evidence that the common eelgrass also harbours unusual telomeres, composed exclusively of the human-type telomere DNA sequence (Figure 1). We have done so by (1) analysis of repeats from NGS data, (2) molecular cytogenetics with telomeric probes using fluorescence *in situ* hybridization (FISH) and (3) identification of its telomerase RNA, including its template region for synthesis of the telomeric motifs. More details on the materials and methods used can be found in the Reporting Summary.

**Figure 1.**
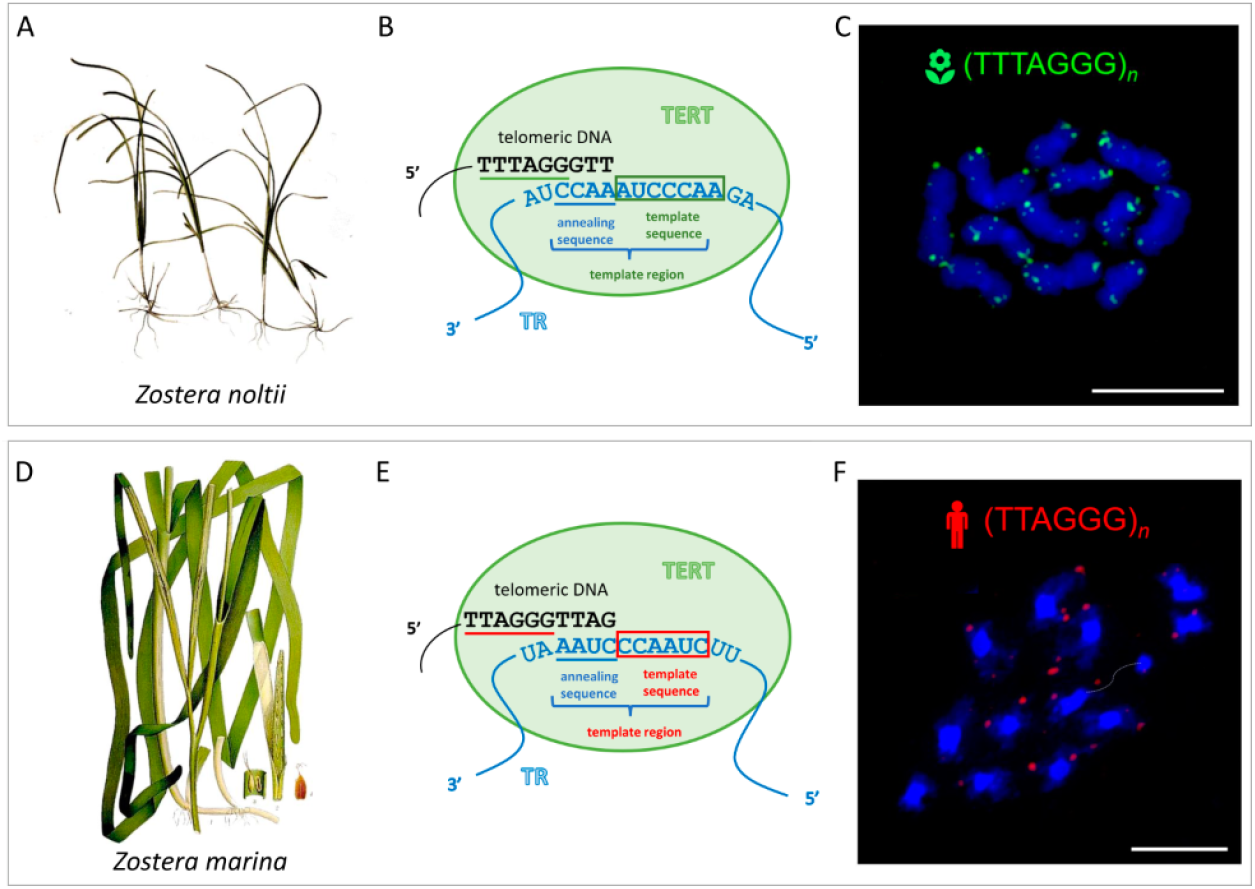
Morphological appearances in *Zostera noltii* and *Z. marina*, respectively - images from Wikipedia - public domain (A, D), schematic model of telomerase RNAs (TR) and telomerase reverse transcriptase (TERT) indicating the annealing and template regions in the synthesis of telomeric DNA (B, E); fluorescence *in situ* hybridization of telomere probes on mitotic chromosomes and (C, F; scale bars = 10 μm).

The first indication that *Z. marina* could have an uncommon telomeric sequence came from the analysis of genomic data from the Finnish population, reported by Olsen et al. (2016), since information about telomere composition was not present in their genomic assembly. The subsequent screening we performed on this dataset revealed the absence of the expected plant-type and the presence of human-type telomere repeats. Although the quality of raw sequence data was excellent, and extreme care was taken to avoid any possible contamination from small marine animals (J. L. Olsen, personal communication), we provide independent evidence of the presence of human-type telomere repeats, as needed to exclude hypothetical contamination and to confirm this exceptional finding.

Thus, in parallel, we sequenced our own materials from Mediterranean populations of *Z. marina* and ran the same genomic screening. We also included the sequencing of a related species, *Z. noltii*, which shares the same habitat. While *Z. marina* genomic telomeric screening yielded exclusively human-type telomere repeats (neglecting reads representing less than 0.001% of the genome portion), *Z. noltii* showed plant-type telomere repeats, also almost exclusively. Based on the total number of reads analysed and the reads containing telomeric sequences, the genome portion of human-type telomeric DNA in *Z. marina* was 0.03-0.3%, while the genome portion of plant-type telomere DNA in *Z. noltii* was 0.06%. The genome proportion of *Z. marina* telomere DNA represented *c*. 3kb of telomere sequence per chromosome arm. This is in accordance with the average telomere length and genome size of *Arabidopsis thaliana*, which is within the known and small genome size range of *Zostera*. However, these are only semi-quantitative estimations, because of the NGS library type, which involves an amplification step biasing the quantitative determination. For more details, see Extended Data (File 1) in the online content. This study was complemented by the search for telomeric motifs in the NGS datasets available for Alismatales species in public databases. We have included species such as the well-known Neptune grass *Posidonia oceanica* (family Posidoneaceae) and other *Zostera* species. The outcome from the tandem repeat analysis, showing the most abundant tandem repeats in all these species, can also be found in the Extended Data (Table 1) in the online content. In all cases, except *Z. marina*, the search yielded the plant-type motif as the most likely telomeric sequence (0.01-7.31% of the genome portion); in comparison, the human-type sequence always represented a much lower proportion of the genome.

Although the analysis of NGS datasets clearly indicated that the *Z. marina* genome unexpectedly harboured these human-type telomeric repeats, cytogenetic confirmation to prove that these ones actually localize to chromosomal ends was needed. To our surprise, no molecular cytogenetic studies of *Zostera* have been reported. Thus, we prepared chromosomes from root tips and analysed them by FISH. The plant-type and human-type probes were independently hybridized to metaphase plates of *Z. marina* and *Z. noltii* (both 2n = 12). Hybridization of *Z. marina* chromosomes with the plant-type probe did not reveal any signal, while, using the human-type probe, weak but clear signals were observed at most chromosome ends. In *Z. noltii*, the opposite FISH profile was obtained: positive hybridization with the plant-type probe, while negative with the human-type probe (Figure 1). As a control for human-type telomere hybridization in plants, we used chromosomes of *Scilla peruviana* (Asparagales), known to possess human-type telomeres^11^ and whose chromosomes did label with the human-type probe in the same experiment (data not shown). To further characterize the chromosome structure in both *Zostera* species, rDNA-FISH was performed. In both species, a single locus of terminal 35S and interstitial 5S rDNA was identified. However, the two species differed in the position of the rDNA loci on the chromosomes: for *Z. marina*, 35S and 5S rDNA were located on one arm of one chromosome while for *Z. noltii*, they were located on different chromosomes (Figure 2).

**Figure 2.**
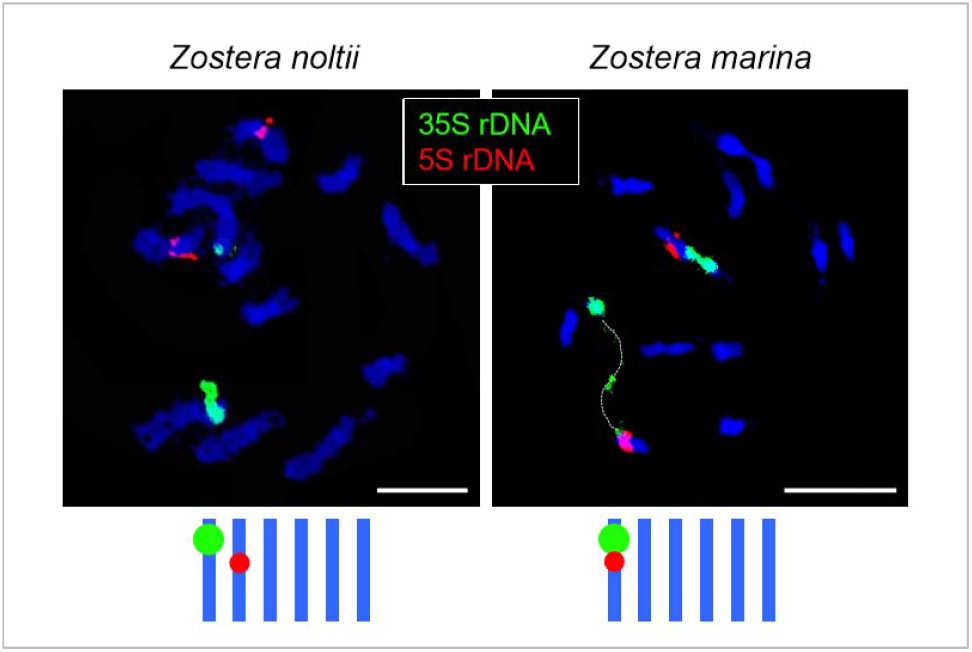
Karyotypes of *Zostera noltii* and *Z. marina*. Both species bear a single terminally located 35S rDNA locus and a single interstitial 5S rDNA locus. While 35S and 5S are situated on different chromosomes in *Z. noltii*, they occupy the same chromosome in *Z. marina*. Note, highly decondensed 35S rDNA site in *Z. marina* (dotted line). The plate of *Z. marina* is incomplete, showing only eleven chromosomes. Scale bars = 10 μm.

## The telomere template in *Z. marina* telomerase RNA helps us understand telomere sequence shifts in closely related species

Finally, we utilized our recent finding of plant telomerase RNA in plants^10^ to obtain independent evidence of the newly found telomeric sequences in *Z. marina*, by analysing its genomic and transcriptomic data. Telomerase RNA is an essential part of the telomerase ribonucleoprotein complex, carrying the template region to produce telomeric DNA by reverse transcription. The template region of the telomerase RNA is usually comprised of two parts. One of them is a complete telomere motif - *“template sequence”* for reverse transcription (Fig. 1C and 1F), and the second is usually only a partial one, which serves as an annealing sequence for the existing telomere DNA. We show here, for the first time, that the telomerase RNA sequences of *Z. marina* and *Z. noltii* serve as human-type and plant-type templates, respectively. The predicted human-type template region of *Z. marina* originated, most likely, from the plant-type template by mutation at its borders. The template region of *Z. marina* could, hypothetically, anneal and synthesize the plant telomere sequence, but the prediction of human sequence annealing and synthesis fits better with the identified telomerase RNA from *Z. marina* and, certainly, with the telomeric repeats detected in this species (see Figures 1 and 3). All other *Zostera* species and Alismatales analysed (from available online datasets) represent exactly the opposite situation, with the plant-type motif in their predicted templates. The reconstruction of the evolution of *Z. marina’s* template region may give us a clue as to the transition from synthesis of the plant-type to the human-type telomere, driven by a more stable mode of annealing of telomere DNA to the changed border of its template region. Complete putative telomerase RNA sequences, in which the corresponding template region is underlined, and other additional results, are presented in Figure 3 and in the Extended Data (Figure 1 and Table 2) in the online content.

**Figure 3.**
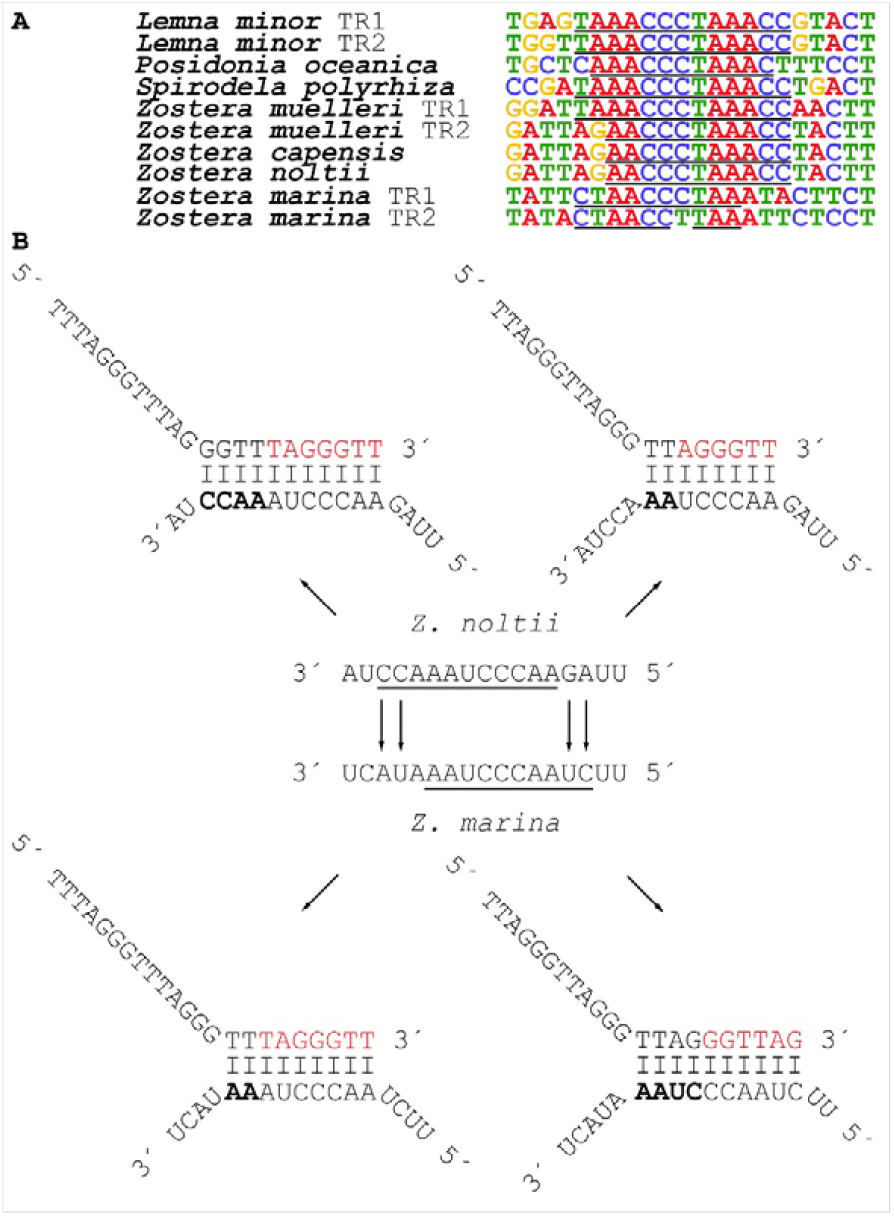
Alignment of telomerase RNAs of several Alismatales species showing the putative template regions (underlined) in which the borders are variable even for plants with the same telomere sequence (e.g. *L. minor, P. oceanica*, and *Z. capensis)* or for paralogs from the same species (e.g., TR1 and TR2 from *Z. muelleri)* (A). The template region may expand, shrink or shift according to the sequence mutation at its borders and adjacent surroundings, as suggested in our model for human- and plant-type telomeric sequence synthesis in *Z. noltii* and *Z. marina*. A more stable mode of annealing of telomere DNA to the changed border of the template region drives the transition from plant-to human-type sequence synthesis in *Z. marina*. (B). The annealing part of template region is highlighted in bold font, the newly synthetized DNA is in red.

Over the last decade, unusual telomeric sequences have been reported for disparate plant groups and in particular, the human-type sequence pops up repeatedly across the green plant phylogeny (Figure 4). Although homoplasy in the evolution of telomere motifs is common, as short/simple motifs like the plant- or the human-type have appeared repeatedly in distant groups^11–14^, it has been hypothesized that the ancestral telomere sequence is the human-type since it is the most commonly found across the tree of life and it is present in the branches close to the possible root of eukaryote phylogeny^15^. In the plant monocot order Asparagales, the human variant occurs along with lower abundances of two or more variants of the minisatellite sequences, including the consensus plant-type telomeric sequence, *Bombyx-type* (TTAGG) and *Tetrahymena-type* (TTGGGG)^11^ as well as the recently described *Allium-type* telomere sequence (CTCGGTTATGGG)^9^. Genus *Genlisea* (eudicots) displays a similar profile: the plant-type is found in some species while others reveal intermingled sequence variants (TTCAGG and TTTCAGG)^16^. In certain groups of algae (Glaucophyta and Chlorophyta), human-type telomeric sequences have already been detected in some species or genera, while others also present the plant-type and other variants^17^. A similar situation applies to the genus *Zostera* in which both human and plant-type telomeric sequences are found, adding another exception to the increasingly questioned conservation of telomeric sequences throughout plants (and perhaps throughout the tree of life).

**Figure 4.**
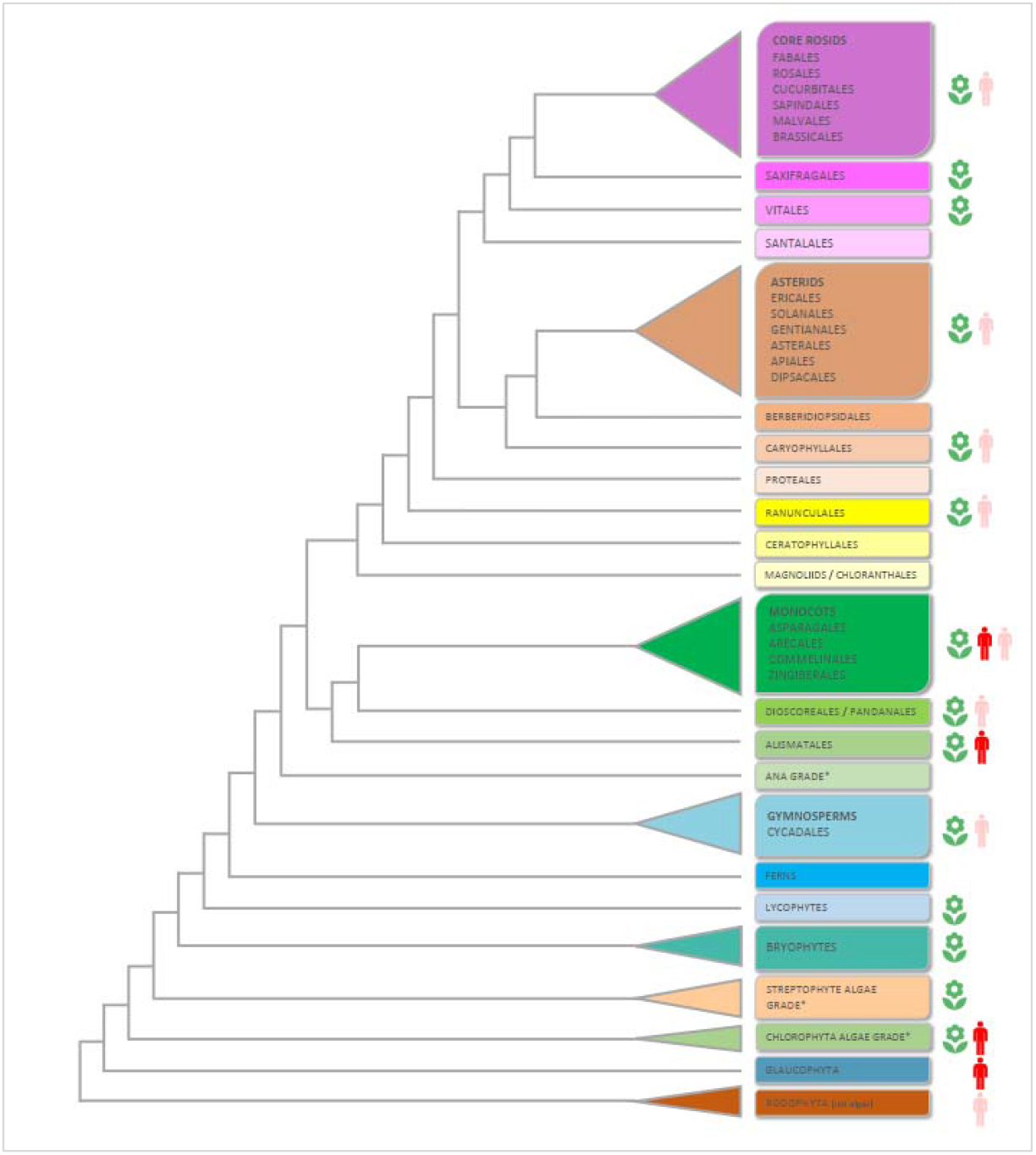
Occurrences of the plant-type and human-type telomere motifs in Archaeplastida (plants in the broad sense), based the APG IV (The Angiosperm Phylogeny Group 2016)^25^ and on the One Thousand Plant Transcriptomes Initiative (2019)^26^ Branch lengths do not express real time scales. For simplicity reasons, some groups have been collapsed or represented by a single branch. Some minor orders (Acorales, Petrosaviales, Trochodendrales, Buxales, Gunnerales, Dilleniales, Zygophyllales, Crossosomatales, Picramniales, Huerteales, Icacinales, Metteniusales, Garryales, Escalloniales, Bruniales and Paracryphilales) not depicted on the tree. The green flower represents the plant-type telomere sequence (TTTAGGG) and the red man the human-type telomere sequence (TTAGGG). The light red man represents cases in which there is some evidence (FISH, slot blot of from NGS data only) that would point to the presence of the human-type sequence, but further work is needed to confirm it.

What are the evolutionary implications of this finding? The cases of telomere shifts in closely related species have huge evolutionary significance, as these not only affect the terminal chromosomal DNA sequence but, if successfully maintained, must be associated with profound changes in the protein components of telomeres, which recognize this telomere DNA in a sequence-specific way. Our data show that the telomere system is more dynamic than previously thought and that evolution tends to converge to a few successful types, since such changes are “evolutionarily demanding”. This may explain the several independent findings of the human-type motif (one of the least complex telomeres) in some plants.

Moreover, a combination of the available genomic data from Alismatales with the recent breakthrough of the discovery of plant telomerase RNA10,18,19 have led us to the astonishing conclusion that the template region in plant telomerase RNA is a very dynamic element, with unexpectedly variable borders, even in species with the same telomere sequence. The finding of two distinct telomerase RNA paralogs within several of the Alismatales genomes analysed, including the *Z. marina* genome assembly, offers an answer to how plants are able to change their telomeres. In the genome assembly of *Z. marina* we found two telomerase RNA gene candidates, however, only one with the human-like template region was detected in the transcriptomic analysis (Extended Data Table 2). Hence, some plants can use mutations in the template regions (particularly at its borders) of the telomerase RNA paralogs to escape from telomere sequence uniformity. Thus, our detailed analysis of the *Zostera marina* genome has revealed the first model for the transition from the plant to the human telomere sequence.

## Online content

Methods along with any additional Extended Data items are available in the online version of the paper; references unique to these sections appear only in the online paper.

## Reporting summary

Further information on research design is available in the Nature Research Reporting Summary linked to this paper.

## Data availability

Raw NGS data files have been deposited in the Sequence Read Archive (SRA) of the National Center for Biotechnology Information (NCBI). The genomic skimming whole-genome shotgun sequences for *Zostera marina* and *Z. noltii* are accessible in bioproject PRJNA594842. The raw transcriptomic data for *Z. marina* are accessible in SRR10664371.

## Acknowledgements

We are thankful to Jeanine Olsen who kindly discussed several aspects on this paper; Javier Romero and Francesca Rossi for their careful indications on where to find the studied *Zostera* and to Jordi Rull, Amelia Gómez and Joan Pere-Pascual Díaz who assisted in plant collection and DNA extraction. This work was supported by: ERDF [project SYMBIT, reg. no. CZ.02.1.01/0.0/0.0/15_003/0000477], Czech Science Foundation [17-09644S and 19-03442S], Ministry of Education, Youth and Sports of the Czech Republic [project CEITEC 2020 (LQ1601)], and by the Spanish [CGL2016-75694-P (AEI/FEDER, UE)] and Catalan [grant number 2017SGR1116] governments. VP benefited from an EMBO Short-Term fellowship (grant no. 7368) and SG is the holder of a Ramón y Cajal contract (RYC-2014-16608). We also acknowledge the CF Genomics CEITEC MU supported by the NCMG research infrastructure (LM2015091 funded by MEYS CR). Access to computing and storage facilities owned by parties and projects contributing to the National Grid Infrastructure MetaCentrum (https://metavo.metacentrum.cz/en/index.html) provided under the programme “Projects of Large Research, Development, and Innovations Infrastructures” (CESNET LM2015042), is greatly appreciated.

## Author Contributions

VP and SG initiated the project. VP, MM and SG performed the bioinformatic search of telomere motifs in SRA datasets. VP, SG and DV obtained and processed *Z. marina* and *Z. noltii* material for DNA extraction and chromosome preparation. TM performed the molecular cytogenetics experiments. PF and JF searched the TR subunit in the NGS datasets. SG and VP wrote the manuscript and all authors participated in the manuscript revision.

## METHODS

### Plant material collection, DNA and RNA extraction

Young plants of *Zostera marina* and *Z. noltii* were collected in the wild (Location: Étang de Thau, beach next to the Musée de l’Étang de Thau, Sète, France. Collectors: Sònia Garcia, Amelia Gómez, Vratislav Peska, Jordi Rull & Daniel Vitales. Date: 20.02.2018). Vouchers are deposited at the Herbarium of the Institut Botanic de Barcelona with the numbers BC973540 (Z. *noltii)* and BC973541 (Z. *marina)*. The leaves were carefully cleaned and inspected for the absence of other organisms. After that, they were wiped by paper tissue and frozen on dry ice. They were stored at −80 °C until usage. Genomic DNA was extracted by a CTAB method^20^ and the quality checked with Qubit Fluorometric Quantification (ThermoFisher Scientific, 128 Waltham, Massachusetts, USA). Actively growing young roots were harvested from collected plants, and subsequently split into two aliquotes. The first aliquote was pre-treated with 0.05% aqueous colchicine for 2h 30 min at room temperature, fixed in ethanol/acetic acid (3:1, v/v) fixative for 24 h at 4 °C and stored at −20 °C until further use. The second aliquote was frozen on dry ice and stored at −80 °C until usage in the RNA extraction by TRI Reagent^®^ (Sigma-Aldrich) according to the manufacturer’s instructions. The extracted RNA was checked on an Agilent 2200 TapeStation system (Agilent Technologies) using RNA ScreenTape^®^ (Agilent Technologies) and RNA concentration was determined by a Qubit 2.0 fluorometer using a Qubit™ RNA BR Assay Kit. Ribosomal RNAs were depleted from 5 μg of total RNA of the sample by Ribo-Zero rRNA Removal Kit (Seed, Root) (Illumina^®^). After rRNA depletion, samples were diluted in 10 μl of RNase-free water and 5 μl of the sample were used for RNA library construction using NEBNext^®^ Ultra™ II Directional RNA Library Prep Kit for Illumina^®^ (NEB) according to the manufacturer’s instructions. Library was treated by a fragment size selection in range from c. 200 to 500 bp using AGENCOURT^®^AMPURE^®^XP magnetic beads (Beckman Coulter). Library quality was checked by an Agilent 2200 TapeStation system using High Sensitivity D1000 ScreenTape^®^ (Agilent Technologies).

### Illumina sequencing

Extracted genomic DNA was sent to the NGS provider (BGI Genomics, Shenzen, China) and skimming genomic sequencing on Hiseq4000 was ordered for each sample. Approximately 15 million of 2 × 150 bp PE reads were obtained from each species. The cDNA Library (total RNA, rRNA depleted, and converted to cDNA) was sequenced in CEITEC MU Genomics Core Facility on a NextSeq500 platform (Illumina^®^) using NextSeq 500 v2.5., mid Output 150 cycles kits (Illumina^®^) for 2 × 75 bp PE reads. Raw reads are deposited in GenBank under the accession numbers SRR10664371 (RNAseq from *Z. marina)*, SRR10664317 (genomic NGS data for *Z. marina)* and SRR10664318 (genomic NGS data for *Z. noltii*).

### NGS data analysis

The telomere sequences from Alismatales were searched in the genomic skimming data obtained from BGI and GenBank by Tandem Repeats Finder (TRFi) tool (https://tandem.bu.edu/trf/trf.html): *Alocasia odora* (Araceae) - SRR7121940, *Halophila ovalis* (Hydrocharitaceae) - SRR5877255, *Lemna gibba* (Araceae) - SRR074103, *Lemna minor* (Araceae) - SRR2882980, *Posidonia oceanica* (Posidoniaceae) - SRR2315671, *Spirodela polyrhiza* (Araceae) - SRR7548932, *Wolffia australiana* (Araceae) - SRX3579183, *Valisneria spinulosa* (Hydrocharitaceae) - SRR6038670, *Zostera muelleri* (Zosteraceae) - SRR1714574, *Zostera marina* (Zosteraceae) - SRR3926352. *Zostera marina* (Zosteraceae) - SRR10664317, our dataset, *Zostera noltii* (Zosteraceae) - SRR10664318, our dataset. Subsequent analysis with custom made scripts were performed according to Peska et al. (2017)^21^. Briefly, the TRFi script was set to look for the short tandems with the unit size of 4-50 nt in at least pentamerous array. Motifs were sorted in descending order in each species. Human and plant telomere motifs were manually found in the final output. A summarized output of the most abundant tandem repeats, including telomere motifs, is presented in the Extended Data (Table 1). RNA- seq *de novo* assembly of data published in GenBank (SRR10664371) was done using Trinity- v2.7.0 (https://github.com/trinityrnaseq/trinityrnaseq/wiki). The assembly was performed as stranded RNA-seq with paired-end fastq datasets. Putative TR from *Z. marina* were found out using blast with published orthologs^10,18,19^.

### Telomerase RNA search analysis

Telomerase RNA (TR) orthologs were identified in publicly available datasets, going from TRs published from Alismatales using novel TR candidates in BLAST searches. The final set of TR presented in this work was aligned in Geneious 8.1.9 (https://www.geneious.com) using MAFFT alignment (Algorithm: E-INS-I; Scoring matrix: 200PAM/k = 2; gap open penalty: 1.53).

### Chromosome preparation

Chromosome spreads from root tips were prepared according to a published protocol^22^. Briefly, selected root tips were rinsed in distilled water (twice for 5 min) and sodium citrate buffer (10 mM sodium citrate, pH 4.8; twice for 5 min), and digested in 0.3% (w/v) cellulase, cytohelicase and pectolyase (all Sigma-Aldrich, St Louis, MO, USA) in citrate buffer at 37 °C for 90 min. After digestion, individual root tips were dissected on a microscope slide in approximately 10 μl acetic acid and covered with a cover slip. The cell material was then spread evenly by tapping, thumb pressing and gentle flame-heating. Finally, the slide was quickly frozen in liquid nitrogen and the cover slip flicked off with a razor blade. The slide was fixed in ethanol-acetic acid (3:1) and air-dried. Ready-to-use chromosome spreads were checked under a phase contrast microscope for suitable mitotic chromosome figures and the amount of cytoplasm. When appropriate, preparations were treated with 100 μg/ml RNase (AppliChem) in 2× sodium saline citrate (SSC; 20× SSC: 3 M sodium chloride, 300 mM trisodium citrate, pH 7.0) for 60 min and 0.1 mg/ml pepsin (Sigma) in 0.01 M HCl at 37 °C for 5 min, then post-fixed in 4% formaldehyde in 2× SSC for 10 min, washed in 2× SSC twice for 5 min, and dehydrated in an ethanol series (70%, 90%, and 100%, 2 min each).

### DNA probes preparation

The plant-type (TTTAGGG)n and human-type (TTAGGG)n telomere repeat probes were prepared according to protocol for non-template PCR^23^. The *Arabidopsis thaliana* BAC clone T15P10 (AF167571) bearing 35S rRNA gene repeats was used for in situ localization of nucleolar organizer regions (NORs), and the *A. thaliana* clone pCT4.2 (M65137), corresponding to a 500 bp 5S rDNA repeat, was used for localization of 5S rDNA loci. All DNA probes were labeled with biotin-or digoxigenin-dUTP by nick translation as described by Mandáková and Lysak (2016b)^24^. Labeled probes were ethanol precipitated, desiccated and dissolved in 20 μl of 50% formamide and 10% dextran-sulfate in 2× SSC for 3 h.

### *In situ* hybridization and microscopy

For FISH, 20 μl of the labeled probe were pipetted on a chromosome-containing spread and immediately denatured on a hot plate at 80 °C for 2 min. Hybridization was carried out in a moist chamber at 37°C overnight, followed by post-hybridization washing in 20% formamide in 2× SSC at 42 °C. The immunodetection of hapten-labeled probes was performed as described by Mandáková and Lysak (2016b)^24^. Chromosomes were counterstained with DAPI (2 μg/ml) in Vectashield. Fluorescence signals were analysed and photographed using a Zeiss Axioimager Z2 epifluorescence microscope and a CoolCube camera (MetaSystems). Images were acquired separately for all three fluorochromes using appropriate excitation and emission filters (AHF Analysentechnik). The monochromatic images were pseudocolored and merged using Photoshop CS software (Adobe Systems).

---

**Extended Data Figure 1.**
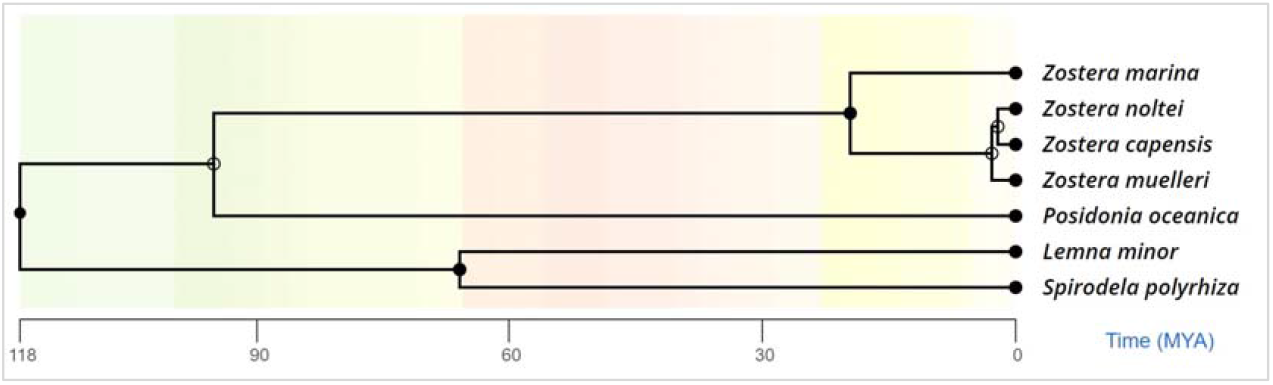
Tree showing phylogenetic relationships between selected Alismatales based on TimeTree.org^27^. *Zostera marina*, which is the only one with human-like telomeres, stands outside of the group formed by the other *Zostera* species analysed in this work.

**Extended Data Table 1.**

Report on the most abundant tandem repeats from TRFi analysis in several Alismatales species. Plant-type telomere sequences indicated in green and human-type telomere sequences indicated in blue.

(see independent file Extended Data Table 1.pdf)

**Extended Data Table 2.**
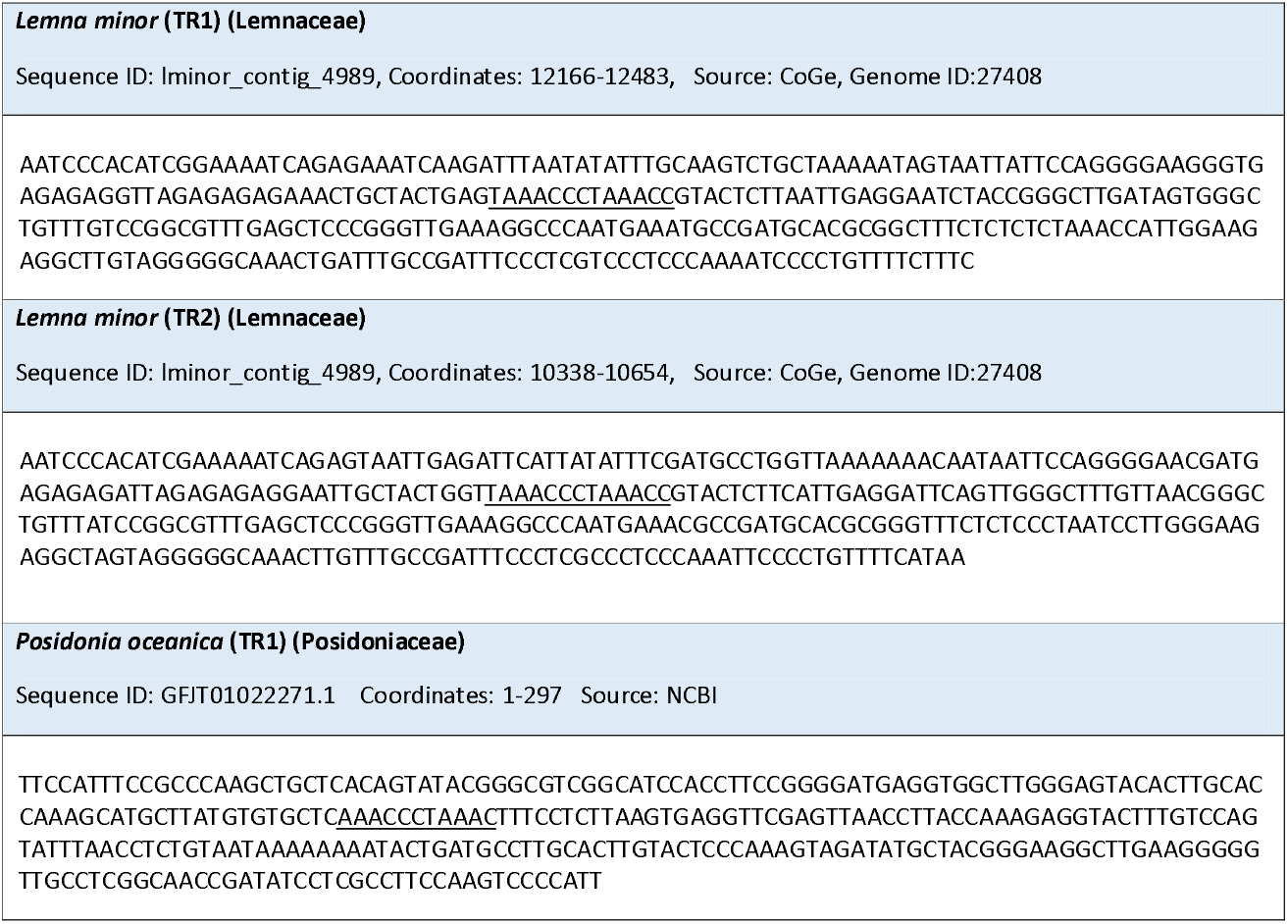
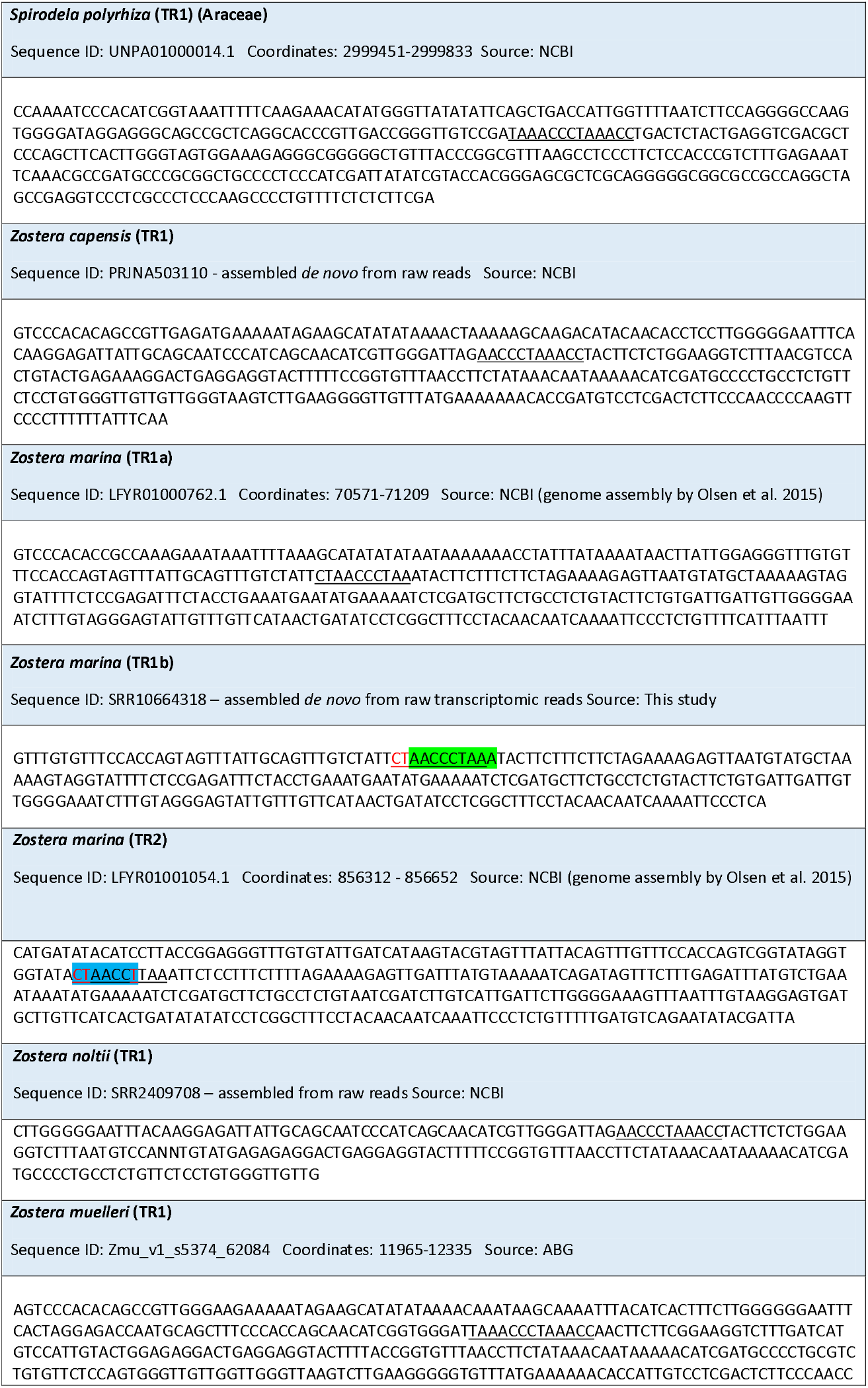
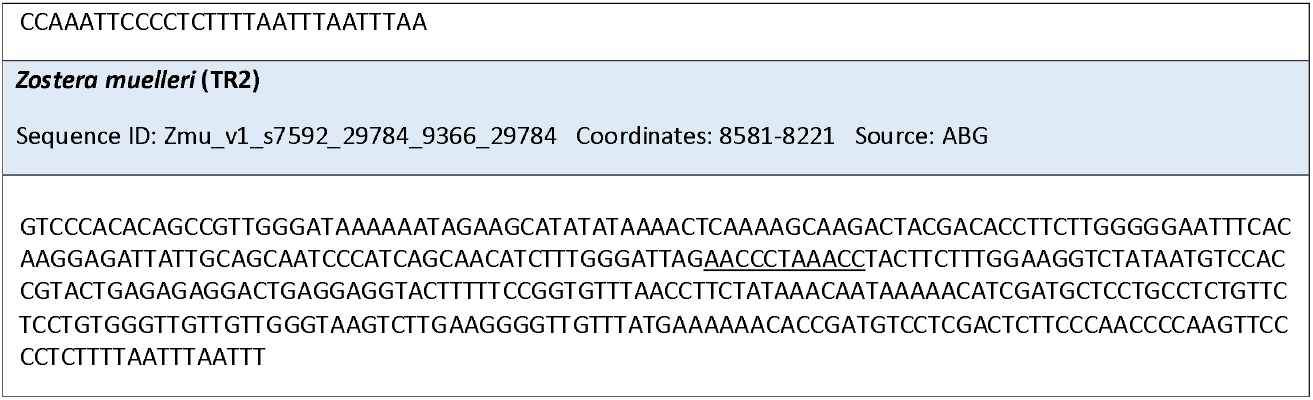
Predicted telomerase RNA sequences of Alismatales species. The sequences of the predicted telomerase RNAs were obtained from or ABG – Applied Bioinformatics Group (http://appliedbioinformatics.com.au/index.php/Seagrass_Zmu_Genome), NCBI -National Center for Biotechnology Information (https://www.ncbi.nlm.nih.gov/), and CoGe – Comparative Genomics (https://genomevolution.org/coge/). TRI and TR2 indicate two possible paralogs for telomerase RNA. TR1a and TR2 from *Z. marina* are genomic sequences. TR1a includes complete promoter and terminator non-transcribed sequences, while TR2 from *Z. marina* lacks several promoter elements. The TR1b from *Z. marina* was assembled for this study from reads obtained in RNA-Seq from root tips. Underlined nucleotides represent the predicted template regions. Note, the red dinucleotide of CT in *Z. marina* TR1b, which as an evolutionary novelty changes the template annealing potential to the preferred human-like telomere sequence. On the other hand, the potential for the plant telomere motif synthesis has not been completely lost in *Z. marina* template motif (green highlighting). In the non-transcribed TR2 of *Z. marina*, there is an additional mutation (C --> T) in the template region, which would potentially lead to (TTAGG)n (known from insects, blue highlighting) telomere sequence. From this point of view, the *Z. marina* TR represents transitional state producing human-type sequence, with a trace of plant-type synthesis.

