## Supplementary figures and images for "Human-like telomeres in *Zostera marina* reveal a mode of transition from the plant to the human telomeric sequences"

### Extended Data Figure 1

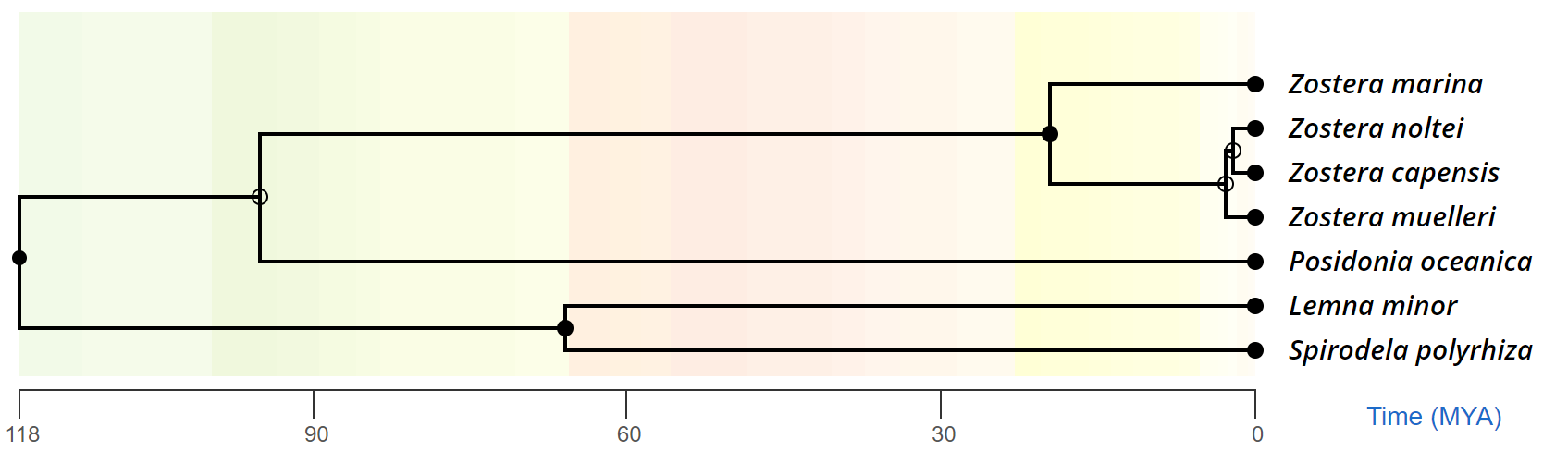
