## Extended Data Table 1 for "Human-like telomeres in *Zostera marina* reveal a mode of transition from the plant to the human telomeric sequences"

| species | *Spirodela polyrhiza* | | *Lemna gibba* | | *Lemna minor* | | *Wolffia australiana* | | *Alocasia odora* | | *Halophila ovalis* | |
|---|---|---|---|---|---|---|---|---|---|---|---|---|
| sample from dataset | SRR7548932 | | SRR074103 | | SRR2882980 | | SRX3579183 | | SRR7121940 | | SRR5877255 | |
| number of analysed reads | | 2000000 | | 2000000 | | 11431597 | | 2000000 | | 2000000 | | 2000000 |
| each species sorted separate | | ↓ | | ↓ | | ↓ | | ↓ | | ↓ | | ↓ |
| *Arabidopsis* motif | CTCT | 33410 | AACCCTA | 146270 | ATAG | 12581 | GAGATA | 82073 | AACCCTA | 696 | ATAG | 32180 |
| human motif | AACCCTA | 28962 | GTAT | 70012 | CTCT | 3678 | AACCCTA | 27114 | GCAGCAACA | 566 | AACCCTA | 17847 |
| | AGAGAG | 15486 | CGTACGTG | 23595 | GCGACTTTAGGA | 2709 | CTCT | 19638 | AAGAA | 556 | TAAA | 3110 |
| | ATAG | 10942 | ACGC | 21681 | AGAGAG | 1737 | ATAG | 13833 | CATATGTCATTTTACACCAA | 479 | ATCGCCTCGATCCCTCG | 2825 |
| | CTCC | 5083 | GCACGTACGCAC | 21056 | AACCCTA | 1323 | AGAGAG | 13779 | AAAAAG | 429 | CTATATCTAATATATTCTAATT | 2600 |
| plant telomere DNA genome portion in % | 1,448 | | 7,314 | | 0,012 | | 1,356 | | 0,035 | | 0,892 | |
| human-like telomere DNA genome portion in % | 0,011 | | 0,0001 | | 0,001 | | 0,048 | | 0,0001 | | 0,012 | |
| | CTCTCTCTCT | 4150 | CACGTACGCACA | 8102 | GTAT | 1118 | GAGAGAGA | 4532 | ATATATCAGTTTGCACCAAC | 368 | GCTCTCAA | 2594 |
| | GAGAGAGA | 4116 | ACAC | 8072 | AGAGGTGG | 1053 | AGAAGA | 4474 | ACCAACATATATCAGTTTAC | 347 | ATCCCTGGATCGTCTCG | 2498 |
| | CTCCCT | 3911 | CTCT | 7130 | AGAGAGAT | 933 | CTCCCT | 3330 | TCAT | 335 | GAATCTT | 2183 |
| | CTCTCTCC | 2470 | CACACA | 6567 | CTTCC | 639 | CTCTCTCC | 3305 | AAACA | 330 | ATCAAGAAA | 2018 |
| | AATATAAATAATAAT | 2157 | AGAGAG | 5895 | GAGAGAGA | 568 | CTCTCTCTCT | 3230 | AAAAAC | 328 | ATCGAGGGATCGAGACG | 1302 |
| | AATAATAA | 2120 | ACATGCAT | 3932 | GAGATA | 542 | ACCCG | 2752 | TAAA | 250 | ACGATGATG | 1238 |
| | TAAA | 1934 | CATGCGTG | 3347 | CTCTCTCTCT | 425 | CTCTTTCTCT | 2401 | CAACATATATCATTTTACAC | 240 | CTATCTAC | 1236 |
| | ACAG | 1833 | ATAG | 2310 | CATC | 402 | AAGAGAGA | 2378 | GTAAGTAT | 235 | ATAGATGG | 1066 |
| | AATATATTAT | 1726 | GATGAT | 2102 | CACAGGC | 400 | AAGAAATC | 2288 | AAATA | 227 | GTAT | 1042 |
| | TATATTAAAATAT | 1514 | CACACACACA | 1930 | AAGAGG | 371 | AGAGAGAT | 1605 | CTTT | 213 | ACACATAC | 996 |
| | CTCTCTCTCC | 1393 | ACGCACATACGC | 1756 | ACAG | 348 | CACCTCA | 1506 | ACTGACCAGATGATCCGATT | 192 | AGTGAGG | 985 |
| | AATAATAATATT | 1352 | CGTGCATGCGTA | 1738 | ATAGAGAGAG | 343 | GTGAA | 1379 | AAACC | 189 | ATACAT | 979 |
| | GAGACA | 1149 | CATC | 1735 | CTTT | 287 | ATAGAGAGAG | 1343 | CCAGCCGTCGG | 174 | AACCCTG | 836 |
| | GTAT | 1115 | CATGCGTACGTA | 1706 | CTCTCTCC | 286 | ATCAATATCGATATC | 1300 | GATCCAATTACTGACCAGAT | 173 | ATCCCTCGATCGCACCG | 789 |
| | AAGAGG | 1062 | CGCACGTATGCA | 1622 | TAAGA | 286 | CTCTCTCTCC | 1287 | ATAT | 168 | ATAT | 788 |
| | GTGA | 945 | CACGTATG | 1547 | CTCCCT | 278 | AGAGATAGAGAT | 1242 | CCGATT | 152 | AGAAAG | 784 |
| | AAATA | 874 | CTCTCTCTCT | 1515 | CGCTCCTAAAGTCGCTC | 269 | AGAAAG | 1232 | AAGAGAATG | 151 | CCCTCGAT | 755 |
| | CTTT | 822 | CACGCACG | 1406 | AGCGTGAGCGACTTTAGGAGTG/ | 264 | AAGAAGAAG | 1224 | CGGCTGGGCGA | 150 | AGATAGAT | 726 |
| | ATTATTAT | 790 | CGCACACA | 1392 | GGCGCTACTTTA | 227 | CCCTAA | 955 | ACCAACATATGTCAGTTTGC | 148 | CATATATA | 714 |
| | GAGAGAGAGAGAGA | 784 | GAGAGAGA | 1324 | AGATAGAT | 222 | AAGAGG | 917 | CAGCCT | 145 | ATACTAT | 709 |
| | GAGATA | 652 | CACGTTCGCACA | 1311 | CTCCCG | 221 | AGAGAGAGA | 848 | GAAACC | 145 | TGCGCGAACAGCGCGAAAC | 698 |
| | ACAGAGAG | 629 | CACGCA | 1285 | CTTTGA | 194 | TAGAAA | 827 | ATATAT | 129 | TCCTAAA | 684 |
| | AGAGAGAT | 610 | ACCACGTACGC | 1238 | GCCCAGGCCA | 191 | CTCC | 792 | ATGTTGGTGCAAAATGACAT | 114 | AGGGCTT | 681 |
| | ATAATATAATAA | 601 | CATGCGTA | 1156 | AGTCAAAGTCAACCCTATC | 190 | GTAT | 781 | GAGGA | 111 | CATC | 603 |
| | GGTTCTAAAAATAGAA | 547 | CACGTTCGCACG | 1151 | CACCTCTT | 185 | CTTT | 775 | ATAG | 110 | ACCTATCC | 602 |
| | GAGAGAGAGAGG | 536 | ACGCACGTACAC | 1143 | CCGGCCCAGT | 167 | AAGGAG | 762 | CAGCGGCTGGG | 104 | AGCTTGGG | 599 |
| | CGAG | 524 | ACACACAC | 1060 | CTCTCTCTCC | 163 | CACGGAATACGAAAAGC | 719 | CATTTTGCACCAACATATAT | 102 | AACGCGGCCGCAT | 594 |
| | ATATATAATATAA | 506 | ACATGCATACAT | 1028 | CTCC | 157 | AGAGAGAGAGAA | 712 | ATTACTGACCAGAACCG | 102 | ATGGACGG | 544 |
| | TAAAATAAA | 496 | CTCTCTCC | 979 | AAGAGAGA | 152 | GAAGA | 709 | ATAAAA | 100 | CCCTAAT | 514 |

| | | | | | | | | | | | |
|---|---|---|---|---|---|---|---|---|---|---|---|
| CATC | 482 | ATGATGGCAC | 849 | GAGACA | 148 | CATC | 679 | CCAGCCGCTGG | 100 | ACCTAA | 485 |
| CCGCCA | 454 | ATACATAC | 774 | AGAGAGAGA | 133 | AGATAGAT | 670 | CAGCAG | 90 | CTAATC | 450 |
| AGAGAGAGAGAG | 450 | CACAAGTATGCA | 747 | CTCTTTCTCT | 131 | AAGATGAATGAAG | 627 | AAAAAAC | 89 | TAGAATATATTAGATA | 426 |
| AAAAATAGAAAGTTCT | 442 | GCAT | 722 | AGAAGA | 118 | ACAGAGAG | 618 | GCAACA | 88 | CCTAATCCCCAT | 425 |
| CTCTCTCG | 413 | GTACGCATGCAT | 717 | GAAGA | 116 | GAGACA | 596 | GAAATGT | 87 | CGCGCAGTTT | 415 |
| CTCTCTC | 402 | CGCACCTA | 711 | AAGAGAG | 116 | CAGCCG | 595 | CAAA | 85 | ATATAT | 411 |
| ATATATT | 393 | ACGTGCCT | 665 | ACAGAGAG | 111 | GAGAGAGAGAGG | 588 | AAAAACAA | 77 | TAAATTTTCATGTAAATTGCT | 392 |
| AGAAGA | 390 | GATGATGAT | 594 | GATGAT | 110 | AAGAGAG | 579 | AGAAAAAA | 76 | AGGGATCGAGACGACCC | 388 |
| CCCCCT | 388 | CACACACACC | 591 | TAAA | 109 | ACAG | 568 | CTCT | 74 | ACCTAC | 379 |
| ATAAAA | 385 | CACCCACA | 581 | CGGCGGCGGCCGCTGGAAGG | 106 | CTCTCTCA | 507 | GTCAA | 74 | ATCCATCTATCT | 368 |
| GAGGA | 370 | CGTACGTACGCA | 578 | AGTCGCTCCGCTCCCTTATAAAT | 103 | AGAGAGAGAGAG | 498 | AAAAAAG | 73 | TCAGGGC | 350 |
| AGAAAG | 349 | TACG | 573 | AAGAAGAAG | 99 | CGATG | 491 | AAGAGG | 69 | ATCGCGGCGACGGGGGG | 337 |
| CTTC | 348 | CTCCCT | 561 | TCCATC | 98 | GAGAGAGAGAGAGA | 462 | GAAGA | 61 | ATACAC | 336 |
| CTCCCCCC | 316 | CTCTCTCTCC | 472 | AGAACA | 98 | CCCATCTCGTTCTTGATCC | 448 | ATATATATAT | 60 | TAGATTAGATAGA | 333 |
| GAAGA | 314 | CGTATGTA | 460 | AGATTGAGAG | 94 | CTCTGTCTCT | 433 | AGTGTA | 59 | ATATAGAT | 322 |
| AGAACA | 309 | ATGATGATGACG | 451 | GAGAGAGAGAGG | 86 | AGAGAGAGAGAT | 376 | GTAT | 53 | TAAAACC | 320 |
| ACGGCG | 301 | ACCTACAC | 442 | AGAGAGAGAGAT | 81 | GATGAT | 370 | AAAAAAT | 52 | ATATATAA | 317 |
| AATAT | 300 | CCTATGCACGCA | 439 | GAGGA | 80 | ATATCCG | 370 | AGAGAG | 51 | ATCATCGTA | 307 |
| ATTA | 294 | CACACC | 434 | AGGCC | 75 | AGAGGAAGA | 365 | AAAAAGAAA | 51 | CTTCCTATCCAT | 299 |
| ATATAATATAA | 279 | TCACCATCA | 404 | AGAAAG | 72 | AATGAAAAAGATG | 365 | AATAT | 51 | AGGCA | 294 |
| AAGGAG | 275 | ATCACC | 402 | GTGA | 72 | CCGTCA | 364 | CTCTCTC | 49 | TCCCCGTCCTAA | 290 |
| AGAGCG | 275 | ACACACGCAC | 376 | CCCTAA | 71 | CTCTCTC | 350 | AAAATC | 49 | CTATCCGATGTCATGC | 283 |
| GCGAGAGGGAGAGAACG | 271 | ACACATAC | 353 | AGGCAG | 70 | TCCATC | 339 | AAAGAAAAAG | 48 | AAGAA | 274 |
| ATAGAGAGAG | 264 | GCAGGCATGTAC | 345 | AAGAA | 68 | AAGAGAGAGAGAGA | 339 | CATATATA | 48 | TAAGCAAGTTACATGAAAAT | 273 |
| AAGAA | 258 | GATGAC | 343 | ACTCTCTCT | 68 | GAGAGAGAGAGAGG | 319 | AAAACAAAA | 45 | ATCCGTCT | 273 |
| CTTCC | 258 | ACGCACACACAT | 334 | GAGAGAGAGAGAGA | 63 | ATGATGG | 318 | TCAA | 44 | AACCGCCACGA | 272 |
| AGAGAGAGAGAT | 244 | TATGTGCGTGCA | 321 | CCGCCA | 61 | AGAGAGGAG | 317 | ATACAT | 42 | AATTTAAGCAAATTACATGA | 263 |
| GAGAGAGAGAGAGG | 237 | ACGATGATG | 305 | GATGATGAT | 60 | ACGTTCTTGATCCCTTCTT | 306 | AAGGAG | 41 | AAATA | 258 |
| ACAC | 226 | CGTGCGTAGGGG | 303 | AAATA | 59 | AGATAGAGATAGAGATAG | 297 | AGAGAGATG | 41 | CTCCCT | 254 |
| CTCCC | 223 | CGCACACGTA | 278 | GATGAC | 58 | GGAGAAGAA | 288 | AATTTTTCTG | 40 | ATATAA | 254 |
| CGTCTT | 221 | TGATGATAA | 277 | ATCACC | 57 | AATTTTTCTG | 283 | CTCCCCCC | 39 | ACCCTAG | 253 |
| CTCTCTCTCG | 217 | GGGTACGT | 275 | CTCTCTCA | 55 | ACACCAT | 283 | ATATAA | 38 | CCCTAA | 244 |
| TCAA | 217 | ATGTATGTATGT | 246 | ATAGATGG | 54 | ATATAG | 266 | CCGCCCC | 37 | CTCGATCGC | 241 |
| AGCAGG | 217 | CATACACACATA | 244 | AGAGAGGAG | 53 | GATGAC | 262 | CTCTCAT | 37 | AGATAGATAGGA | 234 |
| AACCCTG | 216 | ATGAGGATG | 239 | ATACAC | 50 | CACTCACTCTCTCTCG | 258 | AGAGAGAGA | 36 | AGGAGGAGC | 233 |
| CCCTAA | 215 | CGTGCGTACATG | 238 | AGAGAGAGAGAG | 48 | AGATAGATAG | 234 | CGAGGG | 36 | AAAAAG | 223 |
| AAGATG | 215 | ATGGGTGC | 225 | ATACATAC | 48 | ATAGAGATAAAG | 225 | CTCTCTCC | 35 | CTACCTATCTAT | 223 |
| GCCCCC | 203 | ATGATC | 215 | ACATA | 48 | CCCCCT | 220 | CCTCCA | 33 | AGGGAGCA | 222 |
| AAATAATAATAT | 191 | CACACACACACG | 211 | AACCCTG | 48 | AAAGAGAGAA | 219 | AAGAGAGA | 32 | ATGCACAC | 222 |
| CTCCCCCT | 185 | GCGCACGACCT | 211 | CTCTCTC | 47 | AGAAAATCTTGTCAG | 215 | CCCCCT | 32 | AGAGAGAT | 210 |
| CCCCCCT | 183 | AGGAGGAGGACG | 206 | ACATAG | 47 | AAAATTCTAG | 205 | ATAAAAAA | 32 | CGATCATCTCGATCCCT | 210 |
| AGAGAGGAG | 180 | GAGAGAGAGAGG | 203 | AAAAAG | 46 | GAAGAGAGAGA | 201 | CAATCA | 31 | ATACATATAT | 208 |
| AATTT | 180 | CACGAACA | 203 | AAGGAG | 45 | AAAGAAAGAGAG | 201 | GAAAT | 30 | AGGCATGACATCAGAT | 206 |
| GAGAGAGGGG | 176 | GCACACACGTAT | 203 | CTCTGTCTCT | 44 | GCCGCCCAAG | 200 | CTCCCT | 29 | AACCTAAACCCT | 195 |
| CACACA | 174 | CACGCACCTACG | 199 | CCCCCCCCG | 42 | AGAACA | 199 | CCCCCCT | 29 | GCTTTAG | 190 |
| CCTCCA | 173 | GATGAG | 185 | GAGAGAGTGA | 40 | GTAAGGT | 196 | AATGAT | 29 | CAGAATATTC | 189 |
| GAGGGAGAGG | 172 | ATGATCATC | 179 | GAGAGAGAGAGAGAGA | 40 | ACAATTCGC | 196 | CCCCCCCA | 29 | GATAGATAGATA | 186 |
| TCAT | 172 | CACACACACACACG | 170 | AAGATG | 40 | ATACACTTCACC | 193 | AAAGAAAAAA | 28 | AAGAGG | 182 |
| CCCCCCCCT | 170 | CACGTCCGCACG | 166 | GATAGATAGATA | 39 | CTCTCTCG | 190 | ATATATAT | 28 | AGAAGC | 181 |
| ATAT | 170 | ATGTAACATACATGCA | 164 | GCCCCC | 39 | AGAGAGAGACAG | 189 | AAGAGAG | 27 | GGAGAAGAA | 180 |

| | | | | | | | | | | | |
|---|---|---|---|---|---|---|---|---|---|---|---|
| AAGAGAG | 160 | CTCTCTCG | 159 | ACGATGATG | 39 | GAGAGAAGGA | 188 | ATATATT | 27 | GTAG | 179 |
| CCGCCCC | 160 | CTCTCTCTCG | 154 | AGATAGAC | 38 | AGAAAAGAAGA | 188 | CTCC | 26 | AGGGATCGAGATGATCC | 178 |
| CCCCCCCG | 157 | GGAGGA | 152 | AAAAAGAAA | 38 | AGATAGAGAGAT | 185 | ACCCA | 26 | TATACATATAAATA | 177 |
| GATGAT | 156 | TAGATACA | 152 | AGAGAGAGAGAA | 37 | CTTCCTT | 183 | CCCCCCCG | 25 | GAACATTCCA | 177 |
| CTCTGTCTCT | 155 | AGTACGTG | 146 | TCAA | 37 | AAGAAGAAGAAGAAG | 180 | ATACAC | 24 | CGGCAAGGCAA | 177 |
| AGAGAGAGAGAGAGAGAG | 154 | AATGAT | 145 | CCCCCCCG | 37 | CTCTCTCTCG | 179 | AAGAAAAGA | 24 | CCATCCTAATCA | 176 |
| TCCATC | 153 | ATCGTTATC | 145 | CCCCCT | 36 | GAGAGAGTGA | 177 | ATATATATA | 24 | ATATACTT | 174 |
| CATCC | 152 | ACAGACAT | 135 | GAAGAGAGAGA | 36 | GGGAC | 176 | GCCCCC | 23 | CTCTCTC | 172 |
| AGAGGGAGGGAG | 148 | ATCACCATCATC | 134 | AGAAAAAA | 34 | CTCCCCCC | 172 | AGAAGA | 22 | AAGGGAGCTTG | 172 |
| AGGCAG | 144 | ACAGAGAG | 132 | CTCTCTCTCCT | 33 | AGAGAGGA | 170 | CCCCCCCCT | 21 | AAACCCC | 167 |
| AAAAAG | 143 | GCACAACGCAC | 131 | CTCTCTCG | 32 | AAGTGTG | 166 | GAAAAAGAAAA | 21 | CAATCA | 163 |
| ATGAG | 143 | CATGTACGCACA | 130 | GAGAGAGAGAGAGG | 31 | AGGCAGAT | 163 | GAGAGAGA | 20 | GAGGA | 162 |
| CTCCCCCCCC | 138 | GTATGGTAT | 128 | CTCCCCCC | 31 | AAGAA | 159 | GAAGAGAGAGA | 20 | GCGCGAACA | 159 |
| CGGGC | 138 | AATGAC | 122 | AAGAAAG | 30 | AAAAAG | 159 | AGAACA | 20 | CCTCGATCCCTCGATC | 158 |
| CCAGC | 136 | GCACACACGTCT | 119 | AGAGCG | 30 | ACATAG | 158 | AAGAAAG | 20 | AAGGAG | 156 |
| CTCTCTCA | 134 | CTCC | 118 | ATTA | 30 | CTCTCTCTCCT | 155 | CTCCCCCCCC | 20 | CCGCGATACCCCCGTCG | 156 |
| CTCAG | 134 | ATGATGATGACA | 118 | CGTCTT | 28 | GAGGAAGAGA | 152 | ATACATATAT | 20 | ACGGTCGAGGGATCGAG | 154 |
| CCCCCCCCG | 133 | GTGA | 117 | TAGATACA | 28 | ATAGATGG | 152 | CACATT | 20 | AAGAGAG | 153 |
| GGGGAA | 130 | ATGATGAAC | 116 | AGATAGAGAGAT | 27 | AGAGGAAGAAGA | 152 | AAGAAGAA | 19 | AGATGGAC | 153 |
| CCTCCG | 126 | ACAG | 114 | AGAGAGAGAGAGAGAGAG | 27 | GAGAGGGAGAGAGA | 151 | AAATGC | 19 | ATCCCCCGATCGTCTCG | 153 |
| AACTG | 125 | CGAGGA | 113 | ATATATT | 27 | AAACAC | 147 | CAAAAAAAT | 19 | AAACCC | 151 |
| TAAATATTATATA | 124 | ACGGCG | 113 | AGAGAGAGACAG | 26 | CATGGAATACGAAAAGC | 147 | CTTCCTT | 18 | TATACATATAAA | 149 |
| GAGAGAGAGAGAGAGA | 120 | CCTCCTCGT | 112 | ATCTCTCTCTC | 26 | CTCCCCCT | 144 | AAAAAAAAAGA | 18 | ATCGTGGGATCGAGACG | 146 |
| GGAC | 120 | ACGTACATAC | 110 | CCCCCCCCCG | 26 | GAAGAAGAAGAA | 144 | ATATATAA | 18 | ATATAG | 145 |
| ATATAT | 120 | ATACATGCATGC | 107 | AGAGAGAATG | 25 | GATAGATAGATA | 144 | CAAAAAAAAA | 18 | ACCCT | 145 |
| CTCTTTCTCT | 118 | GCACACACGTCC | 105 | AACATG | 25 | CAGCCAGCCGCG | 144 | CTTTCTTTTTT | 18 | ATCGAGGGATCGAGACC | 144 |
| GCAT | 116 | GTGCGTACGTGG | 100 | ACCCG | 24 | CGCACAAATT | 141 | ATATAG | 17 | ATATCAAAA | 143 |
| CAGCAG | 116 | CACGCG | 98 | AGAGGAAGA | 24 | GAGAGAGAGAGAGAGA | 136 | AAATC | 17 | TACACCTACACACC | 143 |
| AGAGAGAGA | 114 | GCATCA | 98 | AAAAAAG | 24 | AGATAC | 136 | CTCTCTCTCC | 16 | CAAGGAT | 137 |
| AGAGAGAGACAG | 114 | AGAGAGAGAGAG | 96 | AAAAAAT | 24 | AAAAAGAAG | 135 | AGAAAG | 16 | AGGAGCA | 134 |
| GAACA | 114 | CGGAC | 95 | CCGCCCC | 24 | AGATAGAGAGAGAG | 134 | AGAGAAGA | 16 | AGTTTCGTGGC | 134 |
| CTCTCTCTCCT | 113 | TCACCACCA | 95 | AGAGAGATG | 23 | AGACGGCCGG | 134 | CCCCCCCCG | 16 | GAGGGATCGAGGCCATC | 133 |
| AGATAGAGAGAT | 109 | ACATAGATACAT | 94 | ACACATAC | 23 | GAAAAGCCACAGAATAC | 131 | ATATATATAG | 16 | AACCCTT | 129 |
| AAAAAAG | 109 | GAGAGAGAGAGA | 91 | ATACAT | 22 | ACCCTAAACCCTAAACCCT | 129 | CTCTCTCTCT | 15 | GAAGA | 123 |
| GAGAGGGAGAGAGAGA | 108 | AAGGAG | 90 | CTCCTA | 22 | AAGATG | 128 | CCCCCCA | 15 | ATAAAA | 123 |
| AGAGAGAGAGAGAGAGAG | 108 | AGATGATGA | 86 | ATGAG | 22 | AACAAGAAG | 124 | AGAGAAA | 15 | ATATATACTAT | 121 |
| ATCACC | 108 | AAGAGAGA | 81 | ATAAAAAA | 20 | AGAGAAAGAA | 123 | AAAAAATAA | 15 | CCCCCCA | 120 |
| AAAAAAT | 105 | CTCTGTCTCT | 77 | CCTCCA | 20 | CTCACACCTCACACCATAC | 122 | AAAATCAA | 15 | ACATCGGATAGACATG | 119 |
| CCCCCCA | 105 | CTCTTTCTCT | 71 | ATATAG | 19 | AGATAGAC | 120 | ACCGA | 15 | AACTGCGCG | 119 |
| AGGC | 102 | AGAGCG | 64 | ACTCTC | 19 | AGAGAGAATG | 119 | AGTA | 15 | CCCCCCCA | 117 |
| ACCCTAG | 102 | GAGACA | 63 | TCAT | 19 | GAGGGAGAGG | 118 | CTCTTTCTCT | 14 | AAGAGAAAGAGG | 117 |
| AATAAAATAATAA | 102 | AGAGAGAGA | 62 | GATGAG | 19 | ATCTCTATCTCTATCTTT | 118 | CTCTCTCA | 14 | ACAGCGCGAA | 116 |
| GGAAGGA | 101 | GAGAGAGAGAGAGG | 62 | ATGATGATGACG | 19 | AGATAGATAGATAGACAG | 116 | TATCA | 14 | AGAGAGAGA | 114 |
| CGAT | 101 | GAGAGAGAGAGC | 49 | CTCTCAT | 19 | CCCCCCT | 114 | AAAAAAGAAGA | 14 | CTACCCAT | 114 |
| AATGAG | 100 | ATAGAGAGAG | 48 | AATAT | 19 | AGAGGGAGGGAG | 114 | CCCCCCCGCCC | 14 | CTCCCTTGCCAAG | 114 |
| GATGAC | 99 | AGAGAGAGAGAA | 46 | AGAACC | 19 | ATTCCGTGGCTTTTCAT | 114 | CCTCCG | 14 | ACATGACTAGAA | 113 |
| CCCCCCCCCG | 98 | AGAGAGGAG | 44 | AGAGATAGAGAT | 18 | CCCGACCCGA | 112 | TCAG | 14 | ATATATAT | 108 |
| CCCCCCCA | 96 | CTCTCTC | 40 | CTCCCCCT | 18 | GAGAGAGAGAAAGAGA | 109 | CAAAATT | 14 | CCCATCCGAATA | 108 |
| CGAA | 96 | GAGAGAGTGA | 39 | AGAGAGAGTGAG | 18 | AGAGAGAGTGAG | 108 | AAAATCA | 14 | AAGGTAGGGGC | 108 |
| AGAGAGAGAGAA | 95 | AGAGAGAT | 37 | GATGAAGAA | 17 | ACTCTC | 107 | GATGAG | 13 | ATATACTAT | 107 |

| | | | | | | | | | | | |
|---|---|---|---|---|---|---|---|---|---|---|---|
| CCAGA | 94 | GAGATA | 36 | GTAG | 17 | CGGAC | 107 | CGATC | 13 | ATGGATAGATAA | 107 |
| ACTCTC | 92 | CCAC | 33 | ACCACG | 17 | AGATAGATATAG | 107 | AAAGAAAAGAAA | 13 | CGAAATTCGAC | 107 |
| TAAATAA | 92 | ACATAG | 32 | ATGATC | 17 | ACACGA | 106 | AGTGAGG | 13 | ATATATT | 106 |
| AGCCGA | 92 | CTCCA | 28 | AAAAAATAA | 17 | GAGAGAGGGG | 103 | AAAAAGAAG | 12 | ATCTATCTATCTTCCT | 106 |
| ATCTCG | 90 | ATAGATGG | 27 | CTTTC | 17 | AGATTG | 103 | AAACCCC | 12 | ATCTAATCT | 105 |
| ATGGC | 90 | AAGATG | 27 | AACAAGAAG | 16 | CGGATGA | 103 | CCCCTCCCCCC | 12 | GATGATGATTAT | 104 |
| CTCCA | 89 | AGAGGGAGGGAG | 27 | CTACCT | 16 | CTTCGTCTT | 99 | CCCCCCCCA | 12 | CTTT | 103 |
| AGCGGC | 88 | AGAAGA | 26 | CCATA | 16 | AAGAAAG | 99 | CCCCCCCCCCCG | 12 | CCCCCCCCA | 102 |
| TCAG | 88 | CTCTCTCTCCT | 26 | AAAAAAC | 16 | GAGGA | 97 | AACCTC | 12 | CCTAAACCCTAAAT | 102 |
| ATATATAAAT | 88 | TCAT | 26 | AAGAGAAAAA | 16 | GTATTT | 96 | AGAAGAGAA | 11 | ATATATATAG | 102 |
| ACCCCC | 85 | ACCCCC | 25 | ATTCC | 16 | AAAAGAGAGAG | 95 | ACCCCC | 11 | AGAGCG | 100 |
| AAGGCC | 85 | AAGAGAG | 24 | GTCAA | 16 | ATCTTT | 95 | AACACC | 11 | CAAAACC | 99 |
| GGGCA | 85 | GAGGGAGAGG | 24 | AAGAGAGAGAGAGA | 15 | AGAGAAGA | 93 | AGAGAGGAG | 10 | AGGGATC | 99 |
| CTTCCTT | 84 | CCCCCCCA | 23 | AGAGGGAGGGAG | 15 | GAGAGAGAAAAA | 93 | CTCCC | 10 | CTCT | 98 |
| ATATAGAT | 84 | CTCTCTCTCGCTCT | 22 | CGAGGA | 15 | AAGAAGAAT | 89 | CCCCCCCCCA | 10 | AGAGAGAAGA | 96 |
| GAGAGAGTGA | 83 | CATATATA | 22 | CCCCCCCCT | 15 | ATCTCTCTCTC | 86 | CCCCTCT | 9 | AAAGGG | 96 |
| AGCCAC | 83 | AAGAGG | 21 | ATAAAA | 15 | AGAAGAGAA | 86 | AAGAAGGAAG | 9 | CCCCCT | 95 |
| CTCGG | 82 | GTAG | 21 | ACCCCC | 15 | AGAAGAAGAGA | 86 | CCCCCCCCCG | 9 | ACCCCC | 95 |
| ACGC | 81 | CTCTCTCA | 19 | AGCAGG | 15 | CCTCTCTCCCTCTC | 85 | ACGC | 9 | CCTCCTCCA | 93 |
| ACCAGC | 81 | AGAGAGAGTGAG | 19 | CAAAACA | 15 | CCCCTCT | 84 | CCGCCA | 9 | TAATAT | 93 |
| GGAGGA | 80 | CACAGGC | 19 | CTGGGC | 15 | AGAGAGAGAGACAG | 82 | AAAAAATAAA | 9 | ATAAAAAA | 92 |
| ATTATT | 80 | AGAAAG | 18 | ACTA | 15 | AAGAAGAA | 81 | AATGAA | 8 | CCCCCCCCT | 90 |
| CCGGCT | 79 | CTAAT | 18 | CAGCCG | 14 | AGTGGAG | 81 | ACAC | 8 | AGAAAAAA | 90 |
| GCCA | 79 | CATCCATG | 18 | AAACAC | 14 | AGAGAGAGAAGAG | 80 | CTTC | 8 | AATAT | 90 |
| GATGGG | 79 | CCCCCT | 17 | AGATAGAGAGAGAG | 14 | GGGGAA | 79 | AGAAGAAAAA | 8 | CCTCCA | 89 |
| AAGAAGAAG | 78 | CTCCCCCT | 17 | GAGGGAGAGG | 14 | ATACAT | 79 | AGCGGC | 8 | CTCTCCTC | 88 |
| CTCTCCCTCTCC | 78 | ATACAC | 17 | AGAGAGAAGA | 14 | ATATGA | 79 | AAGAGAAAAAAA | 8 | CCCCCCT | 85 |
| CCTCCCTC | 78 | AACCCTAA | 17 | AAACCCC | 14 | AAATA | 78 | AAATGG | 8 | CCATCTATCCAT | 85 |
| CATCGC | 77 | CCCCCCCCA | 17 | ACATGCAT | 14 | AGAGAGAGGAAG | 78 | ATATATATATGT | 8 | ATATATATATGT | 85 |
| CTGGGC | 77 | CTTT | 16 | CCGCC | 14 | AGATAGAGAGAGAT | 78 | AAAAGAGAGAG | 7 | GATGAT | 84 |
| AGAACC | 76 | CCATA | 16 | AACCCTAA | 14 | GATGATGAT | 78 | AGAAGAAGAGA | 7 | CCCCCCCCCA | 84 |
| ATAAAAAA | 74 | GAAGAGAGAGA | 15 | CCCCCCCGCCC | 14 | CGAGGA | 76 | AACATC | 7 | AAAAAAT | 82 |
| AAGAGAGA | 73 | GAGAGAGAGGGG | 15 | ATGATGA | 14 | CTACCT | 76 | AGAGAAAA | 7 | CTCCCCCC | 81 |
| CCCCCCCGCCC | 73 | CCCCCCCG | 15 | AATTT | 14 | AGAAAAGAG | 76 | ATATAGAT | 7 | AAAAAAG | 81 |
| CCCCTCCCCCC | 72 | AAAAAC | 15 | CCGGCT | 14 | CTCCTA | 76 | ATTA | 7 | TCAT | 80 |
| CCCCCCCCA | 72 | CCCCCCA | 15 | GCAT | 14 | AGAGAGAAGA | 74 | ACCCC | 7 | GTATTA | 79 |
| GAGAGAGAGAGC | 71 | CGCC | 15 | GAGAGGGAGAGAGAGA | 13 | CCCCCCCCT | 73 | TAAATAA | 7 | ACAACAC | 77 |
| AGAGAGGGGAG | 71 | GAAGA | 14 | TGATGATAA | 13 | GATTAT | 73 | ATTCA | 7 | CATCCATG | 77 |
| GAGAGAGAGGGG | 69 | AGAGAGAGAGAT | 14 | CAATTT | 13 | AGAGAGAGAGAGAGAGA[illegible] | 72 | AATGAG | 7 | CTCC | 75 |
| CCGCC | 69 | CTCCCCCC | 14 | AGATAGATAG | 12 | ATCCATCTATCT | 72 | AGGGAGCA | 7 | ACATGCAT | 75 |
| AAAATT | 68 | AAACCCC | 14 | CTCTCTCTCG | 12 | AGAAGGAG | 72 | ATTAAAA | 7 | AGAAAGAT | 75 |
| AGGGGAGAG | 67 | AGAGAGAGACAG | 13 | GAAGAAGAAGAA | 12 | GGGAGAGAGAGAGAGAG | 70 | GAAAAAAAGAAA | 7 | CGCACACA | 75 |
| CTCTTG | 66 | CAAA | 13 | ACCCTAAACCCTAAACCCTAA | 12 | AAGAAGAAGGAG | 69 | CACGCG | 7 | AGAAAGAGATAG | 71 |
| GATTAT | 65 | CCCCCCCCCA | 13 | CCCCCCT | 12 | CAAA | 69 | AGAGAGAGAGAA | 6 | ACCTACAC | 68 |
| AGAAAAAA | 65 | AAGAGAGAGAGAGA | 12 | GATTAT | 12 | AAGAAGGAAG | 69 | AGAAAAGAG | 6 | CTCTTTCTCT | 66 |
| AACATG | 65 | AAAAAG | 12 | CAAA | 12 | ATAAAA | 68 | GAGAGAGAGAGC | 6 | AGAGAGGAG | 66 |
| AAAAAGAAA | 64 | GAGAGAGGGG | 12 | CTCCCCCCCC | 12 | TCAA | 68 | GTGA | 6 | AGGCAGAT | 65 |
| CGATC | 64 | GAGGA | 12 | GAGAGAGAGAGC | 12 | GAGAGAGAGGGG | 68 | CACACA | 6 | CTCCCCCCCC | 63 |
| GGAGGGA | 64 | CCTCTCTCCCTCTC | 12 | CCCCTCCCCCC | 12 | AACATC | 67 | CTTTGA | 6 | GAGATA | 61 |
| CATCGG | 64 | ATGATGA | 12 | AGGGGGGGGGGG | 12 | AATGAA | 67 | ACGGCG | 6 | CTCCC | 59 |

| | | | | | | | | | | | |
|---|---|---|---|---|---|---|---|---|---|---|---|
| GATCT | 63 | AAAAAAC | 11 | AAAGAAAAAA | 12 | CTCCCCCCCC | 66 | AGGC | 6 | ACACTAA | 59 |
| ATGATC | 63 | AGATAGAT | 10 | CCCCCCCCA | 12 | AGAGAGATG | 66 | AAAATT | 6 | ATTA | 58 |
| AGATAGATAG | 62 | CCCCCCT | 10 | CCTCCG | 12 | AGAGAGAGGGAGGG | 65 | AAGAGAAAAA | 6 | CCTAAACCCTAAAC | 58 |
| AGAGAGAGGGAGGG | 62 | AGAGAGAGAGACAG | 10 | CGAT | 12 | AATGAT | 65 | AGAGAAAAAAA | 6 | AAATGG | 58 |
| GAAGAGAGAGA | 61 | CCTCCG | 10 | ATAT | 12 | GAGAGAGAGAGC | 64 | AAAAAATTA | 6 | AGCTTGGAG | 58 |
| AGGGGGGGGGGG | 61 | ATTCC | 10 | AGTGAGG | 12 | AGAGAGATGG | 64 | AAACAAAAAAA | 6 | CAAGC | 57 |
| AAATC | 61 | ATACAT | 9 | GAGAGAAGGA | 11 | ATACAC | 64 | TAATAT | 6 | ATTGAAG | 56 |
| GTAG | 61 | ATGAG | 9 | AGAGAGAGAAGAG | 11 | TCAT | 62 | CAGCCG | 5 | AAAGGA | 56 |
| CGCC | 61 | AAAAACAA | 9 | AATGAT | 11 | AAAAAAG | 62 | TCCATC | 5 | AAAATCCAA | 55 |
| AGAAGC | 59 | CCCCACCCCCC | 9 | AGAGAGATGG | 11 | ATCACC | 62 | AAACAC | 5 | AGAGAG | 54 |
| AGATAGAT | 58 | GCCCCC | 8 | TATCA | 11 | AGAAAAAA | 62 | AAGATG | 5 | AGATAGAC | 54 |
| GAAGAGG | 58 | AGAGAGGGGAG | 8 | TCACCATCA | 11 | AATGAC | 60 | CGAGGA | 5 | CCCCACCCCCC | 53 |
| CCCCCCCCCA | 58 | CCCCCCCCG | 8 | CTATCTAC | 11 | TAAA | 58 | AAGAAGAAGGAG | 5 | TAACCCC | 52 |
| CATATATA | 58 | CCGCCCC | 8 | CTCCC | 11 | AGAGGAGAGAGAG | 58 | ATCACC | 5 | AGGGTTG | 51 |
| CCCCCCTCTC | 57 | CTTTGA | 8 | CAAAACC | 11 | AGAGCG | 57 | AGAAAGAAGA | 5 | AAAAAGAAA | 50 |
| GCAACA | 56 | ACCCTAG | 8 | CCCCCCCA | 11 | CTCTCTCTCGCTCT | 57 | GAGAAGAG | 5 | ACAC | 50 |
| ATATATAA | 56 | AGATAGAC | 7 | AGCCAG | 11 | AGATAGATAGAA | 57 | AGAAGC | 5 | AACCTTA | 50 |
| GGGAGAGAGAGAGAGAGA | 55 | AGAGAGAGGAAG | 7 | GGAGAAGAA | 10 | GCCCCC | 56 | AGAGAAAAA | 5 | ATATATATAT | 50 |
| CCGCCGCCA | 55 | AGAGAGATGG | 7 | AACATC | 10 | GGAC | 56 | AAAATTAA | 5 | ATTATTAT | 49 |
| AAAAAATAA | 54 | AGGGGAGAG | 7 | AAACA | 10 | GGAAGGA | 56 | CACACACACA | 5 | ACCCC | 48 |
| GGAGAAGAA | 52 | AAAGCA | 7 | AAAAAAAAAGA | 10 | GAAGAAGAAGAAGAG | 56 | AGCAC | 5 | ATATATATA | 48 |
| CCCCTCT | 52 | CCCCCCCCCCCA | 7 | CCCCACCCCCC | 10 | AGAGAGGGGAG | 55 | ATATATAAAT | 5 | CCCCTCCCCCC | 47 |
| AATGAT | 52 | ACTCTCTCT | 7 | CCTAAACCCTAAAC | 10 | TAACCCC | 55 | CCTGAG | 5 | AAGAAATC | 46 |
| TAGATACA | 52 | AAAGAGAGAA | 6 | AAACCC | 10 | CCCCCCCCG | 54 | AATTAAAT | 5 | GAGAAGAG | 46 |
| CAATC | 51 | ATCTCTCTCTC | 6 | GAGGAAGAGA | 9 | CTCTCCCTCTCC | 54 | CAAAAATC | 5 | CCAAATC | 46 |
| AGAGGAAGA | 50 | AGAAAAGAG | 6 | ATCTCG | 9 | AAAGAAGAAGAA | 54 | CATC | 4 | CTCTCTCC | 45 |
| ATATAA | 49 | CTATCTAC | 6 | ATGATGATGACA | 9 | CCCCCCCG | 52 | CTCCCCCT | 4 | AGAGGAAGA | 45 |
| TATCA | 48 | CAAAACC | 6 | CGAG | 9 | AAAAAAT | 52 | ACTCTC | 4 | ACCCTAAACCCTAAACCCTAA | 45 |
| AGCAC | 48 | CCATCTATCCAT | 6 | CCCCCCCCCA | 9 | AAGAAGAGAGAG | 52 | ATCTCTCTCTC | 4 | CTTCC | 45 |
| AGAGGAGAGAGAG | 47 | AGGGCTT | 6 | CATCGC | 9 | ACGATGATG | 52 | AGAGAGAAGA | 4 | AAAATTAA | 45 |
| AAACA | 47 | CGAG | 6 | ATATAT | 9 | AAACCCC | 51 | CCGCC | 4 | AGATAGATAGAA | 44 |
| CAGCCG | 46 | ATGCACAC | 6 | GAACCC | 9 | TATCA | 50 | AAAGAAAG | 4 | GAAGAGAGAGA | 43 |
| CTTCGTCTT | 46 | GGAGAAGAA | 5 | CAAAATT | 9 | TGATGATAA | 50 | GAAGAGGAG | 4 | CCCAAC | 43 |
| AAAGAAAAAA | 45 | CTTCCTT | 5 | CGGATGA | 8 | CCCCCCCCCG | 49 | CCCCCCCCCCCA | 4 | CAAAAAAAT | 43 |
| AACATC | 44 | CCCCCCCCT | 5 | GGGAGAGAGAGAGAGAGA | 8 | ACATGCAT | 49 | ATTCC | 4 | CCTAAACCCTAAA | 43 |
| AAAAAAC | 44 | ATCCATCTATCT | 5 | GGAC | 8 | CAAGAT | 49 | CGGGC | 4 | CCCTAATCCAAA | 43 |
| GCAGCAACA | 44 | CTCCCCCCCC | 5 | CCCCCCCCCCCG | 8 | AGAAAGAAGA | 48 | CCAAGC | 4 | CTCCTA | 42 |
| CGAGGA | 43 | AGAGAGATG | 5 | CCCCCCA | 8 | AGGGGAGAG | 45 | CAAAAC | 4 | GGAGGA | 42 |
| AAAAAATAAA | 41 | AAAAAAG | 5 | AAAAACAA | 8 | AAAAAGAAA | 45 | GAGACA | 3 | AGAGAAGA | 41 |
| CCCCCCCCCCCA | 40 | AGAGGAGAGAGAG | 5 | CTCTTG | 8 | GTGA | 44 | GGAGAAGAA | 3 | ATCCAA | 41 |
| AACAAGAAG | 39 | CCAGA | 5 | AAACC | 8 | CGTCTT | 44 | GATGAC | 3 | CCCCCCCCG | 40 |
| CCCCACCCCCC | 39 | AGGGTTG | 5 | AATGAG | 8 | ACGC | 44 | CTCTCTCG | 3 | GAAAT | 39 |
| CTTTC | 39 | AAAACAAAA | 5 | ATGGC | 8 | GAAGAGG | 44 | CTCTCTCTCG | 3 | GCATCA | 38 |
| ATACAT | 38 | GTAAGTAT | 5 | CAAAAAAAAA | 8 | CCGCCCC | 43 | AGGGGGGGGGGG | 3 | CATCGC | 37 |
| ATGTATG | 38 | TCCATC | 4 | CTCGG | 8 | CCGCC | 43 | AGGCAG | 3 | CTCTCTCTCCT | 36 |
| CCCCCCCCCCCG | 38 | ATGATGG | 4 | AACATA | 8 | GATGAAGAA | 43 | CCCCACCCCCC | 3 | ATACATAC | 36 |
| AGAGAGAGGAAG | 37 | GAGAGAAGGA | 4 | GAGAGAGGGG | 7 | TCAGGG | 42 | AATTT | 3 | CTCTCTCTCT | 35 |
| ACATA | 37 | AGAGAGGA | 4 | CTCTCTCTCGCTCT | 7 | AGATGATGA | 41 | CACAA | 3 | AGGGGGGGGGGG | 35 |
| ACCCT | 37 | ACTCTC | 4 | AGATGATGA | 7 | GAGAAGAG | 41 | CATCGC | 3 | AGAAGA | 34 |
| CTAATC | 36 | AGAGAGAGGGAGGG | 4 | CCCCCCTCTC | 7 | ACAC | 40 | CGAT | 3 | GAGGAAGAGA | 34 |

| | | | | | | | | | | | | |
|---|---|---|---|---|---|---|---|---|---|---|---|---|
| | GAGGAAGAGA | 35 | AAAAAGAAA | 4 | ACCTAA | 7 | CCCCTCCCCCC | 40 | CACGCA | 3 | CCCCCCCCCCCA | 34 |
| | CCTCTCTCCCTCTC | 35 | CTTC | 4 | AAAAAATAAA | 7 | CCCCCCTCTC | 40 | CTTCC | 3 | ACCCG | 31 |
| | AAAAACAA | 35 | CCCCCCCCCCCG | 4 | TAGAAA | 6 | GATGAG | 40 | CACACC | 3 | AGCAGG | 31 |
| | ACCCC | 34 | AACATG | 4 | AAAAAGAAG | 6 | TCACCATCA | 40 | GTATTA | 3 | AAAAAATAA | 31 |
| | AGCCAG | 34 | ACCCC | 4 | GAGAGAGAGAAAGAGA | 6 | AGAAAGAGATAG | 40 | CAAC | 3 | TCCAAC | 31 |
| | CCAC | 34 | AACCCTG | 4 | AGATTG | 6 | AGAAAGAT | 39 | CAAAACA | 3 | CACACA | 30 |
| | AGATTG | 33 | GATGATGATTAT | 4 | AAAAGAGAGAG | 6 | AGGGGGGGGGGG | 38 | AGGCA | 3 | AGCCCTC | 30 |
| | CTATCTAC | 33 | AAACAAAAAAA | 4 | AAGAAGAAT | 6 | ACACATAC | 38 | CACCCACA | 3 | CTCTCTCTCC | 29 |
| | GCCGAC | 33 | CACCTCTT | 4 | CCTCTCTCCCTCTC | 6 | GGAGGA | 36 | ATCCAA | 3 | AGAGAGAGAGAA | 29 |
| | AACCTTA | 33 | AGAGGAAGA | 3 | CTCTCCCTCTCC | 6 | CCTCCCTC | 36 | ATATATACTAT | 3 | CCGCCCC | 29 |
| | CCAAGC | 33 | AAAGAAAGAGAG | 3 | AAAGAAGAAGAA | 6 | ATACATAC | 36 | GTTTTA | 3 | AATTT | 29 |
| | GATCCA | 33 | AGAACA | 3 | CAAGAT | 6 | ATGAG | 35 | AGAGAGAT | 2 | GCAT | 29 |
| | CGGAC | 32 | AAGAA | 3 | ATATAGAT | 6 | TAGATACA | 35 | CCCTAA | 2 | ACCCA | 28 |
| | CTCTCTCTCGCTCT | 32 | GAGGAAGAGA | 3 | CATCC | 6 | AAACA | 35 | GAGAGAGAGAGG | 2 | CCCAATC | 28 |
| | CTCTCAT | 32 | AAACAC | 3 | GCAACA | 6 | CTATCTAC | 35 | CTCTGTCTCT | 2 | AAGAGAGA | 27 |
| | ACCCA | 32 | GATAGATAGATA | 3 | CGGACGA | 6 | AAAGCA | 35 | AGAGGAAGA | 2 | AAGAAGAAGGAG | 27 |
| | CAGCCT | 32 | AGAGAGAGAAGAG | 3 | CGCC | 6 | ATGTATG | 34 | GAGAGAGAGAGAGG | 2 | ATGAG | 27 |
| | AAGAGAGAGAGAGA | 31 | GATTAT | 3 | AAAGAGAGAA | 5 | ACATA | 34 | AGAAAAGAAGA | 2 | AAAGCA | 27 |
| | GGGAC | 31 | CCCCCCCCCG | 3 | AAGAAGAAGAAGAAG | 5 | CTCCC | 33 | GGGAC | 2 | CTCTCTCG | 26 |
| | ATATATATA | 31 | ATAAAAAA | 3 | AGAGAGGA | 5 | CGATC | 33 | AGAGAAAGAA | 2 | AGATAGAGAGAT | 26 |
| | ATATATATAT | 30 | CGAGGG | 3 | AGTGGAG | 5 | CACACA | 30 | AGAGAGAGAAGAG | 2 | CCCCCCCG | 26 |
| | CTACCT | 29 | CCCCCCCGCCC | 3 | ATCCATCTATCT | 5 | ATAAAAAA | 29 | GGGGAA | 2 | TCAGGG | 26 |
| | GATGAG | 29 | CTCTCAT | 3 | AGAGAGAGGGAGGG | 5 | CGTATGTA | 29 | AGAGGAGAGAGAG | 2 | GGAC | 25 |
| | AAAAAC | 29 | AAAAAATAAA | 3 | AGAGAGGGGAG | 5 | CTTC | 28 | AAAGAAGAAGAA | 2 | GGAGGGA | 25 |
| | TAACCCC | 28 | AACCTTA | 3 | GAGAAGAG | 5 | AAAAAC | 28 | CGTCTT | 2 | ATGATGG | 24 |
| | ACACACAC | 28 | GTATTA | 3 | AAAGCA | 5 | ACACACAC | 28 | GGAGGA | 2 | ATCACC | 24 |
| | AGGGTTG | 28 | CAAC | 3 | AAAAAAGAAGAA | 5 | CAAAACC | 28 | ATGAG | 2 | CTTC | 24 |
| | AAAATC | 28 | ATGGACGG | 3 | CATCGG | 5 | AAAAAAGAAGA | 28 | AAAGCA | 2 | AAAAAC | 24 |
| | CCCAAT | 28 | CAAAAAAAAA | 3 | CAATA | 5 | AGAGAAAA | 28 | CAAAACC | 2 | CCTCCG | 24 |
| | ACCCTAAACCCTAAACCCT | 27 | ACACTAA | 3 | ACCGA | 5 | CTTTGA | 28 | AACCCTAA | 2 | CTTCCTT | 23 |
| | AGAGAGATG | 26 | CCAGC | 3 | ATAGA | 5 | ATTGAAG | 27 | CCATA | 2 | ATAGAGAGAG | 22 |
| | AAATGG | 26 | CCAAGC | 3 | ATGGACGG | 5 | CCGCCA | 26 | ACCTAC | 2 | GCCCCC | 22 |
| | CAAC | 25 | AGTGAGG | 3 | AGCCCTC | 5 | CGAGGG | 26 | ATCTCG | 2 | GAAGAGGAG | 22 |
| | AAACAAAAAAA | 25 | CCCTAAT | 3 | AACTC | 5 | AAATC | 25 | GAACA | 2 | CCCCCCCGCCC | 22 |
| | ATCCATCTATCT | 24 | AGAGGTGG | 3 | AAGGCC | 5 | CCGCCGCCA | 25 | AGAAGGAGG | 2 | AAAAAAC | 22 |
| | AAAACAAAA | 24 | CCCTAA | 2 | CCAAGC | 5 | CAATC | 25 | ACCACT | 2 | CACCCACA | 22 |
| | | | | | | | | | | | The table continues with progressively de | |

| *Vallisneria spinulosa* | | *Posidonia oceanica* | | *Zostera muelleri* | | *Zostera noltii* | | *Zostera marina* Finnish | |
|---|---|---|---|---|---|---|---|---|---|
| SRR6038670 | | SRR2315671 | | SRR1714574 | | SRR10664318 | | SRR3926352 | |
| | 2000000 | | 9934246 | | 2000000 | | 31045798 | | 2000000 |
| | ↓ | | ↓ | | ↓ | | ↓ | | ↓ |
| AAGGCC | 11956 | AACCCTA | 88012 | AGTGTA | 3998 | AAACCCT | 18107 | ATTA | 9858 |
| AACCCTA | 2889 | CTTT | 53355 | CTTC | 2778 | GTGCCCTTCGGGGAGTCTGAATAT | 12689 | CCCTAA | 7477 |
| AAACTCTAAATAGAAACCT | 2337 | AAATA | 9948 | AACCCTA | 1822 | AAATTG | 10065 | GTATGAATGGAATGT | 2606 |
| AGGCCAAGGCC | 1799 | ATAAAA | 7235 | ACATTCATTC | 957 | CCGAAGGGCACATATTCCGACTCC | 2205 | CAATTT | 1766 |
| TCCAACAAAC | 1539 | AGAAGA | 4046 | TCAT | 864 | AAATTAGAATTG | 2087 | TAAA | 1748 |
| 0,144 | | 0,886 | | 0,091 | | 0,058 | | 0,0007 | |
| 0,01 | | 0,0002 | | 0,008 | | 0,003 | | 0,374 | |
| GGCCAGGGCAA | 1223 | AAAAAG | 3231 | TAAA | 661 | ATGATTGTGT | 1646 | CTTT | 1608 |
| AAACCCC | 1165 | AAGAA | 3090 | ATAG | 594 | ATATCGACA | 1438 | AAATA | 1267 |
| AAGGCCTGGGC | 1161 | AAAGGA | 2045 | AAATA | 542 | AATG | 1409 | AAAAAG | 1047 |
| AGTACAAT | 1096 | AAAAAAG | 1894 | CTTT | 520 | ATTGGA | 1280 | AATAATT | 818 |
| AGAGTTTAGGATTCTAAAT | 1066 | TTAAAAACAA | 1859 | AGTTGCAAACTTAAGAGA | 488 | CCTAAC | 1041 | ATAG | 759 |
| CTAAACTCTAAATAGAATC | 1014 | AAGAAGAAG | 1716 | ATGAAGAAGAA | 461 | TCATCT | 1014 | TCCAAC | 685 |
| ACTCCGGCGAG | 987 | AGAAAAAA | 1678 | TTCCCTTAAATTTGCAAC | 413 | ATCATCATCATG | 1011 | TATCA | 677 |
| TAAACTCTAAATGGAAACC | 838 | AGGAAGAA | 1364 | ATTCATTCAA | 402 | AATTTGAATG | 875 | TAGTGCA | 505 |
| AGCCGGAGGAATCGAAAGGCC | 614 | AAAAAGAAA | 1121 | ACTTA | 393 | AGATCCAAACACTGAGAATGTGGCTGC | 844 | GTAT | 490 |
| AGTTTAGGATTCCATTTAG | 603 | TCAA | 1115 | AGAAAAAA | 383 | AGGAGGA | 807 | AAACA | 427 |
| ACCGAACCGAATCCATTAATGAA | 598 | AACCCTAA | 1023 | ACCCTAAC | 378 | AAGA | 758 | ATAAAA | 378 |
| AAAAAG | 538 | AAGAAAAGA | 1021 | AGAGAAGA | 376 | AACAAA | 751 | GGAC | 376 |
| GTAT | 488 | CTCT | 978 | ATTA | 360 | AGCTAGGAGAA | 640 | AAAATT | 371 |
| AGAAGA | 460 | CAAAACC | 926 | GAAGA | 343 | GATAGGATCGTC | 604 | TCAT | 354 |
| CCCAAC | 444 | GAGATCATGATGAGAAATTAAGCTAAAGAGGGGGAT | 912 | ACTA | 330 | GAAGGGCACATGTTCAGACTCCCC | 585 | AAGAA | 351 |
| AGCCAG | 440 | CATCATGATCTCATCCCCTCTTTAGCTTAATTTCT | 785 | GTAT | 327 | CCGACT | 538 | TTATAA | 343 |
| GGTTTCGGTCAACCGAAGACCATTC | 375 | AAAGAAAAAA | 762 | CATC | 321 | AAAAATC | 447 | AATAT | 332 |
| CAGGCCAAGGCCATGG | 363 | AATTAAAACA | 632 | AAGAA | 289 | ATAG | 419 | ACTA | 297 |
| AAGTAACTGTA | 356 | AGCAT | 560 | AATTT | 276 | AAATCTA | 398 | CATC | 273 |
| CATC | 347 | ACCTAA | 549 | CAAACTTAAGGGAAGTTG | 269 | AAAATC | 387 | CAAA | 265 |
| CTCTCTC | 347 | AAAAAAAAAGA | 545 | ATAAAA | 261 | AACAATCAAG | 385 | AATCAAAACCCCAA | 257 |
| ACAG | 346 | AAAATCAA | 545 | ATTTGCAACTTCTCTTAA | 245 | CATC | 382 | ACCCA | 238 |
| CTCT | 344 | AAAACAATTAA | 488 | TCAA | 218 | ATAA | 372 | AATTT | 228 |
| AACTGT | 344 | AAAAAGAAG | 475 | AAAAAG | 207 | AATAA | 369 | GTTAACCG | 225 |
| ACCCG | 329 | GATGAAGAA | 474 | AAGATG | 207 | ACCAGCAGC | 368 | TCAA | 222 |
| GCCTTGGCCGTGGCCT | 327 | AGAGGAAGA | 436 | AAGAAGAA | 201 | ATTCATTCAAATATAC | 338 | AAAAAC | 220 |
| CCTAAACTCTATATGGAAT | 324 | CAAAAATC | 435 | ATCATTCAAA | 199 | ACTTCCACCG | 336 | AACATTCAACAAAACT | 200 |
| TCAA | 320 | AAAATCA | 419 | AAACA | 190 | AACCCTG | 328 | AGAAATGGGG | 182 |
| ATAG | 307 | AAACCC | 416 | ACAAATTCAAC | 186 | AATTATAATTGA | 321 | GGAGAAGTTT | 178 |

| | | | | | | | | | |
|---|---|---|---|---|---|---|---|---|---|
| AGAGAG | 306 | CATCCCCCTCTTTAGCTTAACTTCTCATCATGATCT | 405 | AAAAAC | 183 | AATCTAC | 317 | CTCATAAATCCTT | 177 |
| CTGGACTTGCC | 298 | AAGAAGAA | 392 | AATAT | 174 | TCCAC | 307 | GTTTTA | 175 |
| AACTCCT | 296 | AAAGGAAAG | 392 | ATGATGA | 172 | TTAA | 304 | ACCCTAAC | 173 |
| AATTTCCCCGA | 293 | AAGAGG | 375 | AACTCCCATTAAGTTTGT | 167 | ATCATCTGA | 302 | AATTAAAT | 172 |
| TCAGGG | 290 | CATCCCCCTCTTTAGATTAATTTCTCATCATGATCT | 356 | CCCTAA | 166 | AACTG | 297 | CCTCCA | 170 |
| AAGACCATTCGGTTTCGGTCACCG | 277 | TAAAACC | 349 | ATTGTATGAATG | 162 | AGAAA | 284 | TAATTAATAAT | 165 |
| AGCTCCGGCG | 267 | AAGAGAGA | 347 | AAAATC | 156 | ATTG | 281 | AAAGTCGATCCCAGG | 164 |
| GCACTCACTCAAAACT | 263 | AGAGAG | 336 | ACACAAGA | 155 | CATGATCAGAAGCCT | 271 | TAATTAATTAAA | 162 |
| ACCATTAATC | 262 | AAAATCCAA | 331 | AAAATT | 150 | AGGTTG | 269 | AAGTTCT | 156 |
| ATAGAGTTTAGGTTTCTAA | 260 | AAGAAAG | 322 | ATTCC | 149 | AAACAAAACCCTA | 256 | AATCCTTCATA | 147 |
| GGCCACGCCCAAGGCA | 258 | CTTCGTCTT | 313 | CAAA | 148 | AGAAAA | 254 | CGGACGA | 141 |
| CTTT | 256 | CACATT | 308 | CTAAA | 145 | GGGGAGTTTGAATATGTGCCCTTC | 252 | ATCGGAGTATT | 140 |
| AAACTAAC | 255 | TTAAAACAA | 307 | AGAAAAGAAGA | 132 | ATGAATG | 248 | ACAG | 139 |
| CAAGTCCAGGC | 253 | AGAAGGAA | 300 | TCCAAC | 126 | ACTA | 234 | CTAATC | 138 |
| ATGGTGATTG | 250 | CCAAGC | 294 | CAAC | 125 | AATCATTCATAC | 234 | ATATATT | 137 |
| AAGTA | 229 | AAACA | 257 | AAGAGG | 119 | AATTGAATGTTGTTTAGTTG | 228 | ATCGGAGTATCT | 134 |
| CCCATTC | 223 | CTCATCCCCCTCTTTAGCTTAGCTTCTCATCATGAT | 256 | AAACCC | 117 | AAGAGG | 225 | CTAACTT | 129 |
| CTCC | 221 | AAACAAATTAA | 256 | AAAACAGTAAATTA | 112 | ATAC | 223 | ATAACA | 126 |
| AGCAGG | 213 | AAAGAAAG | 254 | AAGAAGAAG | 105 | CACCA | 209 | ATATATAATATAA | 125 |
| AGGCAG | 212 | AAAGAAAGAAGG | 244 | ACAG | 105 | AAATT | 206 | GTGA | 123 |
| CTAAACTCTAAATAGAACC | 211 | CAGCACTTTACTTT | 239 | GTGA | 103 | AAAC | 200 | CCTAAATCAAAACC | 112 |
| CCCTAA | 203 | AGTTTTTTTA | 237 | CTAAT | 103 | CACCAGAGTGCATTCCATAG | 189 | CAAAACA | 108 |
| CTAACAAACT | 198 | AAACCCC | 233 | ATTCA | 103 | GTGA | 180 | ATCGGAGTATTT | 108 |
| ATAGAGTTTAGGTTTCTAT | 198 | GGAGAAAAAGAA | 227 | AACCTC | 103 | AAAATA | 178 | CTTC | 107 |
| ACCGAAGACCATTCGGTCTCGGTC | 189 | AGAAAGAAGAAA | 225 | AAATC | 102 | AATTGAAATTGG | 177 | ACAATATT | 106 |
| CAAGCCA | 187 | AAAAAC | 224 | ATTAGCGACGG | 101 | ATTCCATTCATACAC | 176 | CGCCTATCTTGGGCAAA | 106 |
| CCTTGGCCTTGG | 186 | TTAAAACAAT | 220 | AAAAAAT | 99 | CAAAA | 174 | GAAAT | 104 |
| CCGCCA | 171 | AAAACCAA | 217 | AACTC | 99 | AACATTCAATTCAACTAAAG | 171 | ATTAAAA | 104 |
| GTGA | 170 | AGAAAAAAAAAA | 212 | AGAAAG | 92 | GCAGCACCG | 168 | AATGAA | 101 |
| CTATATGGAAACCTAAACT | 170 | AGAAAGAAGA | 207 | AAAAAGAAG | 91 | TTAAAT | 166 | AATAAAATTAAA | 99 |
| AAGGAG | 168 | ATCTCATCCCCCTCTTTAGCTTAAATTCTCATCATG | 203 | AACCCTG | 90 | AATTGAACTAAACAACATTC | 166 | CGAT | 96 |
| CTAAACTCTATATAGAATC | 166 | AAAGAAAAAG | 201 | ATAAAAAAA | 83 | AGGATCTTCCAA | 165 | AAAAAAC | 92 |
| AACACTAAACCC | 164 | AAAAAACTT | 200 | CCAAATC | 79 | ACTCGACGACTCC | 164 | AAATGC | 84 |
| AGAAAG | 158 | AAAAACCTAGC | 193 | TCAG | 78 | AAAAAAC | 159 | AAAAAAT | 81 |
| ACTTCCTT | 158 | AGAAGAAAAA | 192 | CTTCCTT | 76 | ATGAATGAATGTATATTTGA | 156 | AAGAGG | 79 |
| AAGATG | 157 | TTAAGAAAACAA | 184 | AACATA | 75 | GAAAATGTTAA | 153 | ATTCA | 79 |
| GTCAA | 155 | AAAAACAA | 183 | ATGAG | 72 | ATTTGTTTC | 144 | AAAATC | 77 |
| GGCAAGGCATG | 152 | AAAAAAGAAGA | 180 | AATGAA | 70 | GACCCCATCGAC | 141 | TCAG | 76 |
| CTCAACAAA | 152 | CCTAAACCCTAAAC | 178 | AGAGGAAGA | 69 | ATATC | 136 | CTAAT | 69 |
| CAAA | 151 | GAAAAAGAAAA | 174 | CAATTT | 68 | GGAAGAAGA | 136 | AAAATCA | 67 |
| AGGC | 151 | AGAAGAGAA | 173 | AAAAAAG | 63 | AAAATT | 135 | GAAGA | 66 |
| ACTA | 150 | GTAT | 167 | ATACAT | 61 | CATTCAATTCAACTAAAGAT | 134 | AAAAACAA | 66 |
| GAGAGAGA | 148 | TAAAAAAAACT | 167 | GATGAAGAA | 60 | AATCA | 131 | CAAC | 64 |
| CTATATGATATGAA | 144 | AAGGAG | 164 | TTCAAA | 60 | AAAAAAT | 130 | ATTCC | 63 |
| ACACTAAACCCAACACTATATCCTAA | 144 | AGATCATGATGAGAAGTTAAGCTAAAGAGGGGATG | 159 | ATTAAAA | 60 | CGATGAA | 126 | CTTTC | 62 |
| GAGAGGTTTTCGGTTACCGAAACC | 142 | AAAGAAGAAGAA | 158 | AAACC | 57 | GAAACTCAATCTTTTGT | 126 | ATATAA | 61 |
| AGGAATCGAAAGAGCAGCCGG | 141 | CAATG | 153 | AGAACA | 55 | CAATT | 125 | AACATA | 61 |
| CATTAATCACCACCAATCAC | 138 | CTTCCTT | 151 | CAAAAC | 55 | ACACAATC | 125 | AAACCCC | 60 |
| AGGCC | 136 | AGAGAAAAAAA | 148 | GAACA | 54 | ATCTAAAATCAAA | 123 | GATCT | 57 |
| ATCCTAAACACTAAACCCAACACTAA | 134 | CTTTCTTTTTT | 148 | AGTA | 54 | AAGG | 121 | TAAGA | 57 |

| | | | | | | | | | |
|---|---|---|---|---|---|---|---|---|---|
| TAAA | 131 | AGAAAAGAAGA | 147 | ACATA | 52 | CTTGCCTCCT | 115 | CTCGG | 56 |
| TCAG | 131 | AGAAGAAGATGA | 143 | AAGGAG | 51 | CTCCAC | 114 | CACACA | 54 |
| CGACCT | 128 | CCTAAACCCAAG | 143 | TAAGA | 51 | GAACGGAC | 114 | GATTAT | 53 |
| CCATTCGGTTTCGGTCAACCGAATC | 126 | AAAAACCCTAGG | 141 | ATAT | 50 | GTCTGTAGTA | 110 | CCCAAC | 52 |
| AGAGAGGAG | 123 | AAAAAAC | 138 | AAAAAATAA | 48 | AGTTG | 109 | ACCGA | 50 |
| AAGAA | 116 | AAAGAAAAGAAA | 138 | AACTG | 48 | AAACT | 108 | CCCAATC | 50 |
| AAACCGAACCGAGTCCATTAATG | 116 | CAAAAAAAT | 138 | AAAAAAC | 47 | ATCCCAGATGT | 107 | CAATC | 49 |
| GGAGACTCGCC | 114 | AAACTTAAAAACAATTAAC | 138 | ATCGGAGTATTT | 47 | ACAAGAA | 104 | TATAAAA | 49 |
| ACCGAACTGGGTTCTGAG | 114 | AAGAAAGAAAGG | 137 | CTCT | 46 | ACCATC | 102 | TAGAAA | 48 |
| AACACTATATCCAAAACACTAAACCC | 113 | AATGCAAG | 136 | GATTAT | 45 | ATTTTAATTGTAATGTATG | 102 | AACACC | 48 |
| GCGAACTCCG | 109 | AAAGTGCTGAAAGG | 128 | TATCA | 45 | CGAGGGTCGCCGAGTGGAGC | 98 | ATCCAA | 47 |
| GAGACA | 107 | CCCCCTCTTTAGTTTAATTTCTCATCATGATCTCAT | 125 | AGAAGAAGAGA | 44 | CAAC | 97 | ATACAT | 44 |
| CCCTATT | 107 | AGAAAG | 124 | AAGAAGGAAG | 42 | ACCCTCA | 95 | AAAATTAA | 44 |
| ATGATGGAAG | 106 | AGAGAAAA | 124 | CAATA | 42 | AAACAGA | 94 | AAACCC | 44 |
| CCAGGCCAAGTCCAGGGCAAGG | 106 | TTCAAA | 124 | CAAAACA | 42 | CAGGTCGACGACTGAAGAGGAGGT | 92 | AAATC | 43 |
| CTTTC | 104 | CTTC | 122 | CACAA | 41 | AATCAACAAAAG | 92 | CTAAA | 43 |
| AACCGAAGACCATTCGGTCTCGGTC | 104 | AAAACAAAA | 121 | AAGAAAG | 40 | GGAGTT | 91 | AGAAAG | 42 |
| GCCAAGGCCATGGCAT | 103 | GAAGAAGAAGAA | 120 | AAACAC | 37 | ATCCACTACCTTCACCAT | 91 | CAAGAT | 41 |
| GAATGGGATGAGGAGAAGAACTGG | 102 | AAGAGAAAAA | 120 | AAAATCC | 37 | CCCAAA | 90 | AACTC | 40 |
| ACCGAAACCGAAAGGTTTTCGGTT | 102 | CACTCATCAATCAAT | 119 | GAGGA | 34 | ATTGTA | 90 | GTAG | 39 |
| AGCAC | 101 | GAAAAAAAGAAA | 117 | GTAG | 34 | AATTGAAATTAGAATTGA | 90 | AAGATG | 38 |
| GAATCTT | 100 | TACCAAAAA | 117 | AATAATT | 34 | ATAATTGTATGA | 89 | AAACC | 37 |
| ACCAGC | 99 | AGAGAAAAA | 116 | AGAGAG | 33 | ATAAAAAA | 87 | GCCA | 37 |
| AAAATCC | 99 | AAGAGAAAAAAA | 113 | ATACAC | 33 | ACATTCAT | 87 | AAAATCC | 36 |
| ACAC | 96 | AGAGGAAGAAGA | 109 | ATATAA | 33 | AAGAAGTCTGA | 87 | AGAAGA | 34 |
| ACCCCC | 96 | AAACAAAAAAA | 108 | ATAGA | 33 | ACAG | 85 | CTCC | 34 |
| CACAA | 96 | CCCTAATCCAAA | 108 | AGAAGA | 32 | ACATTCATTC | 85 | AAACAC | 34 |
| ACCACG | 94 | AAGAAGAAGAAGAAG | 105 | TAGAAA | 32 | ACCACTTTGTGGATCT | 85 | ACAC | 34 |
| AAACA | 93 | CAAAAAAAC | 105 | ATAACA | 32 | AACAAT | 84 | AAAACAAAA | 34 |
| AGAAGGAGG | 92 | AGAAGAAGAAGATGA | 105 | GCCA | 32 | ACCTA | 84 | GCAACA | 34 |
| ACCCA | 92 | GTAAAGGCTGAAA | 105 | AAAAAGAAA | 31 | CAGACTCCCCGAAGGGCACATTTT | 84 | AAAGGA | 34 |
| AGGATC | 91 | AAAAAAGAAGAA | 104 | TATAAAA | 31 | GAGAT | 83 | ATATAT | 33 |
| GCCGAC | 88 | AAAAAGAAAAAAA | 104 | CCTCCA | 31 | AAAATCATGCATATAATTGAACTA | 83 | AGCAC | 33 |
| CCTGAG | 88 | CATGCAGAGTCC | 104 | AGAAGAGAA | 30 | GGAGAT | 82 | CCGATT | 33 |
| ACGC | 86 | TAAA | 103 | AATGAT | 30 | AAACC | 79 | CAATA | 32 |
| CTCCCT | 83 | AAAAAAT | 102 | AAAGGA | 29 | GATCCATTGGAA | 78 | ATAGA | 32 |
| CTTC | 80 | CTCTCTCTCT | 100 | AGATTG | 28 | CTAAACCCTAAACCCTAAACC | 78 | ATGAG | 31 |
| AGAACA | 79 | GAAGA | 98 | GTATTT | 28 | AATTGAAATTGA | 78 | CACGCA | 31 |
| ACCACT | 78 | AGAGAAA | 98 | CAATC | 28 | AGGC | 77 | AAATGG | 30 |
| GCCCCC | 77 | GTTTTA | 97 | AAAGAAAAAG | 28 | CGCCT | 76 | GAGGA | 29 |
| AGCCCTC | 77 | AAAGAAAAGA | 96 | ATATATT | 28 | TCAAAT | 75 | GCAT | 29 |
| TCAT | 76 | CAAG | 94 | ATATATATA | 28 | ATTCAATTGAACTAAAGAAC | 74 | ATCACC | 27 |
| ACCGA | 74 | CCTAAACCCTAAA | 92 | CCAGA | 27 | CCGTCGT | 73 | GATGAC | 26 |
| CACATT | 72 | GGAAAAGAA | 91 | AAGAGAG | 26 | ATCAATCAAACATTC | 73 | AGGC | 25 |
| CCTCCA | 70 | CTCCCT | 86 | CTTTGA | 26 | ATTAG | 72 | AGAACC | 25 |
| GCAACA | 68 | AAAGCA | 84 | CATATATA | 26 | AAAAAAAAT | 72 | AGTA | 25 |
| GAAGA | 67 | CAAAAAAAAA | 82 | AAAACAAAA | 26 | TAGTCGTA | 72 | CTCT | 24 |
| GAGGA | 67 | CTCTTTCTCT | 81 | TCCATC | 25 | CTTCTC | 70 | AACATG | 24 |
| CAGCAG | 62 | GAAGAAGAAGAAGAG | 78 | CTCCCT | 24 | CTCAAG | 69 | GAACCC | 24 |
| ATTCA | 61 | CTTTC | 78 | ATCACC | 24 | CCTCGC | 69 | AAAAAAG | 23 |

| | | | | | | | | | |
|---|---|---|---|---|---|---|---|---|---|
| CTCCTA | 60 | AAAGGAAA | 76 | AAAAAATAAA | 24 | ACTG | 67 | GATGAG | 23 |
| GAACCC | 60 | CAAAAC | 76 | AAAGAAAAGAAA | 24 | TCATACATTACAATTAAAAT | 67 | AGAAAAAA | 22 |
| AAATA | 59 | CCCCCCT | 75 | AACAAGAAG | 23 | AAAAAAAC | 66 | CTCCCT | 21 |
| GTAG | 58 | AAATGC | 75 | ACCCA | 23 | GGCACCGGAAGCATCTGTACC | 66 | GAGACA | 21 |
| AGCAGT | 57 | TAGAAA | 73 | TAAATAA | 23 | ACTCCCCGAAGGGCACAGATTCAG | 66 | GTATTA | 21 |
| CCAC | 57 | ATAAAAAA | 71 | CTAATC | 23 | ACATATA | 66 | CAAAAAAAAA | 21 |
| CCTCCG | 56 | AAAAAATTA | 70 | GAAAT | 23 | GCTTCCGT | 65 | ACACGA | 20 |
| CTGGGC | 55 | AAAAAATAA | 69 | GATGAC | 22 | AAATGA | 63 | ATGGC | 20 |
| AAGAAGAAG | 54 | CTCTCTCC | 67 | CGTCTT | 22 | CAAACAACAA | 63 | CAAAATT | 20 |
| CACGCA | 54 | CCCCCT | 67 | AGAAGC | 22 | AACACC | 62 | CTCCTA | 19 |
| AAGAGG | 53 | GGAGAAGAA | 63 | AGCAGG | 22 | GAAGATGAA | 62 | ATGTATG | 19 |
| AAGAGAG | 53 | GCATCA | 61 | AAAGGG | 22 | AATTAAATTCATACATTAC | 62 | ATAAAAAA | 18 |
| ACGGCG | 52 | GAGAGAGA | 60 | CTTTC | 22 | AAG | 62 | CTTTGA | 18 |
| AGCGGC | 51 | GAGGA | 60 | AGAACC | 22 | AGAACC | 61 | AAGTA | 18 |
| GATCCA | 51 | AAGAAGGAAG | 55 | GAATCTT | 22 | CAAATA | 61 | GAATCTT | 18 |
| AAAAAC | 49 | ATATAG | 54 | GAGATA | 21 | CCAGAAGGCCTAGAT | 61 | CCCCCT | 17 |
| CCCCCCCG | 48 | ACACTAA | 52 | AACATC | 21 | ATCATGGCAAGAACATATT | 61 | ACCTAA | 17 |
| CACACA | 48 | AGAGAGAGAGAA | 51 | ATATAT | 21 | AAATAGAGTCATACAAACTAG | 61 | AGAAGC | 17 |
| CTTCC | 48 | CTCCCCCC | 51 | AAGTA | 21 | AATGTCACTTCTATG | 59 | CATCC | 17 |
| TCCTAAA | 48 | CTAATC | 51 | GATGAG | 19 | AAACATA | 59 | TTCAAA | 17 |
| AGAAAAAA | 47 | CTCTCTCTCC | 50 | CCAGC | 19 | AGAAGA | 58 | CCGGCT | 17 |
| CCCCCT | 46 | CCCAAT | 50 | CAAAATT | 19 | AATGAT | 58 | GGAGAAGAA | 16 |
| CTTCCTT | 46 | CCCCCCCCT | 48 | GTTTTA | 19 | ATTCCAATTCCA | 58 | AAGAAAG | 16 |
| AAAAAAG | 46 | AAAAAATAAA | 46 | GATGAT | 18 | CACACA | 57 | ATATGA | 16 |
| CAAC | 46 | GAAATGT | 44 | ACTCTC | 18 | TGAAGAAA | 57 | ACGC | 16 |
| ACAGAGAG | 45 | AGAGAAGA | 43 | AAAGCA | 18 | TCAAACATTCAT | 57 | AGAGAAA | 16 |
| AACACC | 45 | ATTGAAG | 42 | AGAGAAAA | 18 | CAAACTCCCCGGAGGGCACATATT | 57 | ATGATGA | 16 |
| CAGCCG | 44 | AAGATG | 39 | GAGACA | 17 | AAGAG | 56 | CCTCCTCCA | 16 |
| GATGAC | 44 | CCTCCA | 39 | GATCT | 17 | ATACAT | 56 | ATTATTAT | 16 |
| AGAAGC | 44 | AAAATT | 38 | AGGC | 17 | CTGAAAT | 56 | TAATAT | 16 |
| AGCTTGGAG | 44 | ATCCAA | 37 | CGAA | 17 | AACATC | 55 | ATTCATTCAA | 16 |
| CTCCCCCC | 42 | GGAAGGA | 36 | AAAATTAA | 17 | CACCAAA | 55 | ATATAG | 15 |
| CCCGACCCGA | 42 | TAACCCC | 35 | AAAAAATTA | 17 | AATCATTCAA | 55 | ACATA | 15 |
| CCGCCCC | 41 | CATC | 34 | CAAGC | 17 | AGAACTTACCATCTT | 55 | ACCACT | 15 |
| CCCAATC | 41 | AAACC | 33 | ACAC | 16 | AATTAAG | 55 | GAAATGT | 15 |
| CTCTCTCTCT | 40 | CACAA | 32 | CCATA | 16 | TAAAAAACAT | 54 | CTCCA | 15 |
| CAAAATT | 39 | AAAGAAAGAGAG | 28 | GCAACA | 16 | CTCCGTTCATTCCAAGCCGAGCCTC | 54 | ATATAATATAA | 15 |
| AAAAAGAAA | 38 | ACCAGC | 28 | CACATT | 16 | AACATATAAC | 54 | AAACAAAAAAAA | 15 |
| AGGCA | 38 | AGCCCTC | 28 | GGAC | 15 | CTATGCA | 53 | CAAAAC | 15 |
| CCCCCCCCCG | 37 | AAAATCC | 27 | AGCAT | 15 | GTGAATT | 53 | GAACA | 14 |
| CTCCC | 37 | CTCCCCCCCC | 25 | ATATATATAT | 15 | ATTAGA | 52 | AAAAAATAAA | 14 |
| GCCA | 37 | GAAGAGGAG | 24 | ATCCAA | 15 | AACCAA | 52 | AAAATCAA | 14 |
| AACAAGAAG | 36 | AAGAGAG | 23 | ATATGA | 14 | ATGATACCAAAAAATTGGGT | 52 | CAGCCT | 14 |
| CCCCCCT | 36 | CTCC | 22 | AACATG | 14 | AATAAAGAAC | 52 | CAAG | 14 |
| CCGGCT | 36 | GAGAGAGAGAGAGG | 22 | CTCAG | 14 | AAAAAAG | 51 | TCCTAAA | 14 |
| CGAGGG | 35 | ATATAA | 22 | AACACC | 14 | GAGGA | 51 | AACCCTA | 13 |
| ATACAT | 34 | AGAGAAAGAA | 21 | ATTATT | 14 | AAAAAAAAAT | 51 | AAGAAATC | 13 |
| CCCCCCCA | 34 | AACCTTA | 21 | CCTGAG | 14 | AATCACGTGGGCCCCACCCGGA | 51 | CTCCCCCC | 13 |
| AGCCAC | 34 | AGAGAGAGA | 20 | AGGCA | 14 | CAAAAAAAA | 51 | CCCCCCT | 13 |
| CCCCCCCCT | 33 | CAAA | 20 | TCCTAAA | 14 | TCAAACAT | 51 | AGCAGG | 13 |

| | | | | | | | | | |
|---|---|---|---|---|---|---|---|---|---|
| CCCCCCCCG | 33 | TCCTAAA | 20 | ATACATAC | 12 | GCTTGAGGTTGAGGT | 51 | AACCCTG | 13 |
| CCCCCCA | 33 | GAGAGAGAGAGG | 18 | ATATAGAT | 12 | AGGATCTTCGAT | 51 | AGCAGT | 13 |
| GCAT | 33 | AGATTG | 18 | GCAT | 12 | AATTAAGGTTTG | 51 | ACCAGC | 13 |
| CTCTCTCC | 32 | ACCCCC | 18 | GTATTA | 12 | CAATC | 50 | AACTG | 13 |
| CTCCCCCCCC | 32 | AACCTAAACCCT | 18 | CTCCA | 12 | ATAGAA | 50 | CACATT | 13 |
| ATACAC | 31 | AAATGG | 17 | ATTATTAT | 12 | ACCACCATC | 50 | AGAGAG | 12 |
| AGGGTTG | 31 | CCCTAA | 16 | AAATGG | 12 | CATCTAAAGAGATAGCATC | 50 | CGAGGA | 12 |
| AAGAAAG | 30 | CCCCCCCA | 16 | CAAG | 12 | AAAAATT | 49 | AATGAT | 12 |
| GGAC | 30 | ATACAT | 15 | ATACTAT | 12 | GCACCA | 48 | CCGCCCC | 12 |
| CCCCCCCCA | 30 | CGAGGG | 15 | ACGGCG | 11 | ATCAATA | 48 | CCTCCG | 12 |
| CACACACACA | 30 | CCCCCCA | 14 | CATCC | 11 | GATGAG | 48 | ACCACG | 12 |
| ATAT | 30 | CCCCCCCCCA | 14 | CAATCA | 11 | AGAGAG | 48 | TAAATAA | 12 |
| ATAAAA | 29 | AAAATC | 14 | AAAGGAAA | 11 | AATCTA | 48 | CATATATA | 12 |
| CCGCC | 29 | CTTCC | 14 | AAGTTCT | 11 | CATCACGTCTCCTGAGATACTG | 48 | AAAGGG | 12 |
| AGTGGAG | 28 | CCCCCCCCA | 13 | AAGAGAGA | 10 | CTTTC | 47 | AAAAAATTA | 12 |
| GATGAAGAA | 28 | GGAGGGA | 13 | ACGC | 10 | CATAC | 47 | GAGATA | 11 |
| AAAGAAAAAA | 28 | AGCAGT | 13 | AAAGAAAAAA | 10 | AGAAAGA | 47 | CTCTCTCC | 11 |
| CCCCCCCCCA | 28 | CTCTCTC | 12 | AGGCAG | 10 | AGGATCTTCCAATGGATCGTCGAT | 47 | GTATTT | 11 |
| ATTCC | 28 | AAAGAGAGAA | 12 | AAAAACAA | 10 | AATCAAAACCCCAA | 46 | AACATC | 11 |
| GGAGAAGAA | 26 | AGTA | 12 | TACG | 10 | GAGA | 46 | ATACAC | 11 |
| ATAACA | 26 | CCAAATC | 12 | GCATCA | 10 | CACAAA | 46 | CGATC | 11 |
| CGCACACA | 26 | AGAAAAGAG | 11 | AAGAAATC | 9 | CCCCCCAC | 46 | ACCCCC | 11 |
| CAAG | 26 | AAGAAGAAGGAG | 11 | AGAGAGAT | 9 | CCCCACC | 45 | AGAAGGAA | 11 |
| AGAGAGAGA | 25 | CTCCC | 11 | CTCTCTCTCC | 9 | GGCAAGAACAA | 45 | AATAATAA | 11 |
| CGAGGA | 25 | CCCCACCCCCC | 11 | CGATC | 9 | CCTCTCTC | 44 | AGGATC | 11 |
| ATCACC | 25 | CTAAA | 11 | ACATAG | 8 | ATCAATC | 44 | CTCTCTCTCT | 10 |
| ATGATGA | 25 | CCGCCCC | 10 | ATCTTT | 8 | CCTAATTTCCAAC | 44 | AAAAAGAAG | 10 |
| CATATATA | 25 | GATGAG | 10 | CAAGAT | 8 | CATTCAC | 44 | TGATGATAA | 10 |
| GAAACC | 25 | AACCCTG | 10 | CCCCCCA | 8 | ATCGGAGTATTT | 43 | AATGAG | 10 |
| AGGAGCA | 25 | GCAACA | 10 | AAAAAAAAAAGA | 8 | AATAT | 43 | CTTCC | 10 |
| CACACC | 24 | CCCTATT | 10 | AGAGAAAAA | 8 | CGTC | 43 | GATCCA | 10 |
| GGGCA | 24 | AGAGAGGA | 9 | TAAAATAAA | 8 | ATAAAC | 43 | ATATATATAT | 10 |
| CTCTCTCTCCT | 23 | AATGAA | 9 | CAAAAAAAAA | 8 | AGAAC | 43 | CCCCCCCCT | 9 |
| AAACAC | 23 | AACACC | 9 | AGGAAGAA | 8 | CCGTTCG | 43 | GCCCCC | 9 |
| CCCCTCCCCCC | 23 | AAGTA | 9 | AAATGC | 8 | CATTCAAATATA | 43 | CCCCCCCG | 9 |
| AAATC | 23 | ATAG | 8 | CCGATT | 8 | AGAAGCTA | 43 | ACCTAC | 9 |
| CCGCCGCCA | 23 | TCCATC | 8 | AAAATCA | 8 | CTATCTTGGGCAAACTCGC | 42 | AGCCAC | 9 |
| CAAGC | 23 | AAGAGAGAGAGAGA | 8 | AGGCC | 8 | TAAGA | 42 | CGCC | 9 |
| CCCCCCCGCCC | 22 | AAACAC | 8 | TTATAA | 8 | AAACCCC | 42 | AGGCA | 9 |
| AAACC | 22 | CGAGGA | 8 | ATAGATGG | 7 | CAAAGT | 42 | GAGAGAGA | 8 |
| ACAACAC | 22 | ATACAC | 8 | CGAGGA | 7 | ACTT | 41 | AAAGAGAGAA | 8 |
| CTCTGTCTCT | 21 | CCCCCCCCG | 8 | CCGCCA | 7 | ACCCAC | 41 | ACATAG | 8 |
| ACCCTAAACCCTAAACCCTAA | 21 | GGAGGA | 8 | ACCTAA | 7 | CACCGGAAGCATCTGGACCGG | 41 | AGATTG | 8 |
| AGAGAAGA | 21 | CCGCCA | 8 | ACCACT | 7 | AATAAATCAACA | 41 | ACACATAC | 8 |
| ACCCC | 21 | ATTCC | 8 | AGCAC | 7 | AGCC | 40 | GGAGGA | 8 |
| CAATA | 21 | ACAACAC | 8 | GCTCTCAA | 7 | ATGGAA | 40 | CGGGC | 8 |
| ACCCTAAC | 21 | ATAGA | 8 | ATATAG | 6 | CTCCACCAC | 40 | ATATATAA | 8 |
| CTCTCTCTCC | 20 | CCCTAAT | 8 | AAGAAGAAT | 6 | TCAGAAT | 40 | CAAAAAAAC | 8 |
| GATGAG | 20 | GAGAGAGAGAGAGA | 7 | GGGGAA | 6 | TAGATACATG | 40 | TACG | 8 |
| CGAG | 20 | AGAAGAAGAGA | 7 | CTCCTA | 6 | ACATTCATAGAAGTC | 40 | CCAGC | 8 |

| | | | | | | | | | |
|---|---|---|---|---|---|---|---|---|---|
| AAAGAAAAAG | 20 | GAAGAGG | 7 | GCCCCC | 6 | AGAACA | 39 | ATATATATA | 8 |
| CATCGC | 20 | ATTA | 7 | ACACATAC | 6 | AGAATC | 38 | ATATATAT | 8 |
| AAACCC | 20 | AGCAGG | 7 | CAAAACC | 6 | GCATGTGCGC | 38 | ACTTA | 8 |
| AAGAGAGA | 19 | AATAT | 7 | CCTCCG | 6 | CGGCGGCGAAAGAGAGACGGCG | 38 | ACCCG | 7 |
| AACATC | 19 | CCCAAC | 7 | CGAG | 6 | ACATACACA | 38 | AAGGAG | 7 |
| TACG | 19 | CCTCCTCCA | 7 | AGCAGT | 6 | AATGGAAAGAATT | 38 | AGAACA | 7 |
| AAAAAAC | 18 | CAATA | 7 | CGAT | 6 | AAATGAAA | 38 | AGAGCG | 7 |
| ACCTAA | 18 | CAAAATT | 7 | CCCAAT | 6 | AATGA | 37 | GGAAGGA | 7 |
| CATCGG | 18 | ACAG | 6 | AAGAAAAGA | 6 | CAGAAAT | 37 | CCCCCCCCCG | 7 |
| AGAACC | 18 | AGAGAGGAG | 6 | CACACC | 6 | ACTCGACGCCTCC | 37 | CGTCTT | 7 |
| AATAT | 17 | AGAAGGAG | 6 | GAACCC | 6 | AACAC | 36 | CCGCCA | 7 |
| ATAGA | 17 | CCCCTCCCCCC | 6 | AATTAAAT | 6 | CTGCTT | 36 | CCCCCCCA | 7 |
| GATGAT | 16 | AGAAGGAGG | 6 | ATATACTT | 6 | CCCCCCCCA | 36 | AGAAGAAGATGA | 7 |
| TATCA | 16 | ACCCC | 6 | GTGAA | 5 | GCTTCAGGTTGAGGT | 36 | CACAA | 7 |
| CGTCTT | 16 | AGGGCTT | 6 | ACAGAGAG | 5 | CAAAGGTCT | 36 | CACACC | 7 |
| ATAAAAAA | 16 | AAAATTAA | 6 | CGGAC | 5 | GGAGAG | 35 | CCAAGC | 7 |
| ACACACAC | 16 | AAAGGG | 6 | AGATGATGA | 5 | AAAAGG | 35 | CCTGAG | 7 |
| CCCCCCCCCCCG | 16 | CAGCAG | 6 | ATGTATG | 5 | TACGTGACGCTGTATCTCAGGA | 35 | AGAGAGAGAGAG | 6 |
| CAGCCT | 16 | AGAACA | 5 | ACCCCC | 5 | GACGTGATGCTGTATCTCAGGA | 35 | ACTCTC | 6 |
| CCATA | 15 | GAGAGAAGGA | 5 | CTTCC | 5 | CAGTCGATGGCC | 35 | CCCCCCCCG | 6 |
| GGAGGGA | 15 | AGAGAGAGGGAGGG | 5 | CCGGCT | 5 | CAACC | 34 | AAAAAGAAA | 6 |
| CCCCACCCCCC | 15 | TCAT | 5 | AGAAAAAAAAAA | 5 | ATTTGG | 34 | CCCCCCA | 6 |
| ATATAT | 15 | ATCACC | 5 | AATAATAA | 5 | AAAAAAATAAA | 34 | AAAAAAAAAGA | 6 |
| CCTCCTCGT | 15 | CCCCCCCG | 5 | AAAAAGAAAAAAA | 5 | GTGGAA | 34 | AGCCAG | 6 |
| AGAGGAAGA | 14 | CCCCCCCCCCCA | 5 | ACCAGC | 5 | AACCCAA | 34 | CATCGC | 6 |
| GGAGGA | 14 | AACCTC | 5 | CCAAGC | 5 | CATGAATCAAAACCAAA | 34 | AAAAAATAA | 6 |
| ATGGC | 14 | AACTC | 5 | GTCAA | 5 | CAATTCAACTAAAAACATT | 34 | AATATATTAT | 6 |
| TCAGGGC | 14 | ACCCTAG | 5 | TAAAACC | 5 | ACTCCGGCCCTCCCG | 34 | ATTATT | 6 |
| CTCCCCCT | 13 | GAGACA | 4 | TTAAAACAA | 5 | AATTTAAACATCCA | 34 | GGAAAAGAA | 6 |
| CAATC | 13 | CGATG | 4 | AGATAGAT | 4 | TAATTA | 33 | CTCTTTCTCT | 5 |
| CTCCCG | 13 | GAGGGAGAGG | 4 | GGAGAAGAA | 4 | CAGAC | 33 | AAGAGAGA | 5 |
| CCCCCCCCCCCA | 13 | AAAAGAGAGAG | 4 | GAAGAAGAAGAA | 4 | GACGAG | 33 | CTCTCTCG | 5 |
| CGAA | 13 | AACATC | 4 | AGAGAAAGAA | 4 | AAAGGG | 33 | CGGAC | 5 |
| CGGGC | 13 | AGAGCG | 4 | AGTGGAG | 4 | TTCTAAA | 33 | AGAAGAGAA | 5 |
| CAAAACA | 13 | CCCCCCCCCG | 4 | CTACCT | 4 | GAGTCA | 33 | AAGAAGAA | 5 |
| CCAAGC | 13 | ACATA | 4 | AGAAAGAAGA | 4 | GTTCCA | 33 | GGGGAA | 5 |

creasing values - the last row here is the one with the smallest value of vertebrate motif

| | |
|---|---|
| *Zostera marina Mediterranean* | |
| SRR10664317 | |
| | 30798046 |
| | ↓ |
| CCTAAC | 9472 |
| TTAA | 3247 |
| ATTCCATTCATACAC | 2715 |
| AAATTG | 2464 |
| AAGA | 2419 |
| 0,0002 | |
| 0,031 | |
| ATAA | 1756 |
| AATAA | 1509 |
| AGAAAA | 1356 |
| CACCAGAGTGCATTCCATAG | 1190 |
| ATAG | 1076 |
| ATATC | 841 |
| CTATGCA | 828 |
| AATCAAAACCCCAA | 812 |
| GGAGTT | 781 |
| AAAATT | 625 |
| ATAC | 516 |
| AAAATA | 494 |
| ACTA | 491 |
| CATC | 485 |
| AGAAA | 485 |
| CAAAA | 468 |
| GGTTAACC | 465 |
| AATG | 460 |
| AACATTCAACAAAACT | 460 |
| CTATCTTGGGCAAACTCGC | 439 |
| CAACC | 416 |
| ATTG | 392 |
| AAATT | 332 |
| CGCGAGCCATTCCAGGGACT( | 331 |
| AAAC | 313 |
| ATCATAAACCCCAA | 302 |
| GAATTTGAATGATTTTACAG | 298 |
| ATCGGAGTATTT | 291 |
| CCGCCATCTAAAA | 286 |

| | |
|---|---|
| AATTATT | 283 |
| GGGAGAAATG | 282 |
| TAAAAATAAAAT | 282 |
| ATTTATGAGAAGG | 268 |
| AATAT | 260 |
| CGTC | 248 |
| TAATTA | 245 |
| ACAG | 242 |
| AACAAT | 236 |
| AGCTGAGCCTGACAAGCCCT | 236 |
| CTCCAC | 235 |
| GTTTGGAGAA | 235 |
| ACCCCA | 219 |
| AACTG | 217 |
| AGGAAGTCGACCTC | 215 |
| AGGC | 208 |
| AAGG | 200 |
| AAAAAAT | 199 |
| AAGTTCT | 195 |
| AGCTGAGCCG | 189 |
| GTTTTA | 186 |
| AACAAA | 185 |
| CAATGATCCGATAGATACTCC | 181 |
| CTTGGGATTTTTTAAAAAGTC | 163 |
| ATAAAAAA | 161 |
| CCCAACTTCT | 160 |
| ATAAATCCTTC | 159 |
| ACTCTCCCAA | 158 |
| AGCCG | 153 |
| AGAATC | 146 |
| ATAGATCATACAAA | 146 |
| GTGA | 144 |
| CCTAAATCAAAACC | 144 |
| AACCCTG | 143 |
| ATATATTCAACCATTCATCCA | 142 |
| CAATAACA | 141 |
| CAAAACA | 141 |
| ATCGGAGTATCT | 140 |
| GATTAG | 138 |
| AAGTCGATCCCAGGG | 138 |
| TAGTACA | 133 |
| AAAATC | 130 |
| TCATTTATAAGAAA | 130 |
| AAGAG | 127 |
| ACTCGCCTGTCTAGGGGCAA | 127 |
| ATACAT | 126 |
| AGCTGAGCCGAGCCA | 126 |
| ATAGATCATAAAAA | 126 |
| CAATT | 125 |
| CTAGATAGGCGAGTTTGCCC | 125 |

| | |
|---|---|
| ATCGGAGTATT | 122 |
| AAATGA | 120 |
| ATAATAG | 119 |
| AATGA | 118 |
| ATTAG | 115 |
| AAGAGG | 114 |
| ATCATA | 114 |
| TCATCCAAAAATATATTTAAC( | 113 |
| GTTAGAA | 113 |
| GTTTGCCCCAAGATAGGCGA | 112 |
| GATCT | 110 |
| ATTGGGGAAATAAT | 107 |
| CAGGATCAGCAACAACTCCA( | 107 |
| ATTTGC | 104 |
| AAAAATT | 103 |
| ACTGAAATTT | 103 |
| AGAACC | 101 |
| CCCAAA | 100 |
| AGAACGTACTTCAATTGCAGA | 99 |
| AGGTCGACTTCTG | 99 |
| CTTCTAACCCTAA | 99 |
| ATATATCATATCATGTATATG | 98 |
| CTGAATCGACGATTCCCAA | 96 |
| AAAGAAATA | 95 |
| ATAGTGCATAGTTCG | 94 |
| CTAG | 94 |
| AGGATGGACACCTTGCAAG | 94 |
| AAAATCC | 92 |
| AACACC | 91 |
| AATCA | 90 |
| AAATGAATTT | 89 |
| TTAAAAAAACCCTGTAATA | 87 |
| ATTTATGAGAAGA | 86 |
| CCGTCGT | 85 |
| CCACAGCAGGATCAGCAACA/ | 85 |
| GAGAT | 84 |
| AGTGCATAGTACAT | 84 |
| AAAGAGA | 83 |
| ACCATC | 82 |
| ACTG | 82 |
| CAATA | 82 |
| CAGCAACATTTC | 82 |
| ATCGGATCATTG | 81 |
| AACATTCAAAAAAACT | 81 |
| CAAC | 80 |
| ACTCTGATAGTGAGCCAG | 80 |
| ATCATAC | 80 |
| AAAAAAG | 79 |
| AAACACTACAT | 79 |
| AAAAATGAAATT | 79 |

| | |
|---|---|
| CACTAACCATAA | 78 |
| AAAAAAAAT | 77 |
| GAATTGTATT | 77 |
| TGGATCTGGTTCAGGAAG | 76 |
| AAATTCCTGT | 74 |
| CGAT | 73 |
| AAAAAAC | 72 |
| CCATTGTTTCCACCA | 71 |
| AACCCTAAATCAAA | 70 |
| AAAAATC | 69 |
| AGCC | 69 |
| ATGATTA | 69 |
| CATATATATGTACCATAT | 69 |
| CAATC | 68 |
| AACAAAATTAACATTC | 68 |
| ATGAC | 67 |
| ATAGGCGAGTTTGCCCTAG | 67 |
| ATAATAT | 67 |
| ATTTATATTATAT | 67 |
| CACACA | 66 |
| GAGA | 66 |
| CACAAA | 66 |
| ATGGAA | 66 |
| CACTAA | 66 |
| ATGGCATTTCTAAAC | 66 |
| AAACCCT | 65 |
| AATGGAA | 65 |
| AGACA | 64 |
| GAACC | 64 |
| AAACC | 63 |
| CTTTC | 63 |
| ATATGG | 63 |
| CCCTAC | 62 |
| CCGACCTTGATGCCTCGAGG[illegible] | 62 |
| AAGTCGACCTCAT | 62 |
| GAGGA | 61 |
| AAAAAAAAAT | 61 |
| ATAAAC | 61 |
| CAGAC | 61 |
| ATAAACAAT | 60 |
| CCGCCATCTAAAGA | 60 |
| GAATGGGTTT | 60 |
| AAATTGATATTA | 60 |
| ATTTGG | 59 |
| CATCA | 59 |
| CATG | 59 |
| CAAG | 59 |
| AAATCAAACCCCCA | 59 |
| ATGAATGATTGAATATATTTT[illegible] | 58 |
| GGAGAG | 57 |

| | |
|---|---|
| AGCCCG | 57 |
| CATTTTAA | 57 |
| ACATCATCTTCATTATGT | 57 |
| GCACCA | 56 |
| GATCC | 56 |
| AAAATAAC | 56 |
| GATTCCAATTCAATCGCC | 56 |
| ATGTAGGATATGATATGATAT | 56 |
| CATTCCAAAACATACCATTC | 56 |
| CCAATCTCGTTTCCC | 56 |
| AGTTG | 55 |
| AAAAAAAC | 55 |
| CGGTGAGGAGAGGTCCGACG | 55 |
| CGTTCTCTGTAT | 55 |
| TAAGA | 54 |
| AAAAAATC | 53 |
| GTAAGAGGAACAAGTAATTG | 53 |
| AAACT | 52 |
| CCAGT | 52 |
| CCCTT | 52 |
| AGCAC | 52 |
| ATATAG | 52 |
| CAATGGGCAC | 52 |
| ATAGTTC | 52 |
| ACTATTTTGAATT | 52 |
| GTATGATATCATATCATAATA | 52 |
| TCATCT | 51 |
| ACACG | 51 |
| AATTAATTAAAT | 51 |
| CGAA | 50 |
| AGCCTG | 50 |
| TAATTAAA | 50 |
| CCCCCA | 49 |
| CATTC | 49 |
| AAACAGA | 48 |
| AAAAGG | 48 |
| CTTTTTTT | 48 |
| ACGC | 48 |
| CCCGGTTCTGTCGCTCCGGCT | 48 |
| CCACCACCATTA | 48 |
| CAAGAACCCACCACCGGTATA | 48 |
| AACAC | 47 |
| ATCGTCGCTAA | 47 |
| AATATTA | 47 |
| AGGT | 46 |
| AACAT | 46 |
| ATGGTTGA | 46 |
| CCCCCCAC | 45 |
| CCCCCCCT | 44 |
| CCATC | 44 |

| | |
|---|---|
| CAACAG | 44 |
| AGATCA | 44 |
| GAAATTGGCTGATTTTGACTT | 44 |
| ACCACAGTCGAATATACTAGC | 44 |
| CACCA | 43 |
| AAAAAAATAAA | 43 |
| AATATA | 43 |
| AAATCCTAACATTC | 43 |
| TCACTATCAGAGTCTGGT | 43 |
| AAACATACTTCAATTGCAGAG | 43 |
| ATCGGAGTATTG | 43 |
| AAATCACGTGGGCCCCACGG | 42 |
| CCAGAG | 42 |
| CGGAAT | 42 |
| CGATT | 42 |
| CACCTCCTC | 42 |
| ATACCTGAACATGCAACTAGT | 42 |
| AGTAA | 42 |
| AATGCC | 42 |
| AATCACGTGGGCCCCACCCGC | 41 |
| CATGG | 41 |
| AAAAAATT | 41 |
| ATATCTGGA | 41 |
| AATCTTG | 40 |
| ATGTTC | 40 |
| GAGTAAAACGTACTTCAATTG | 40 |
| ATTAATCAA | 40 |
| AGCAACATTTTC | 40 |
| CGAACC | 39 |
| ACCCGACAAGG | 39 |
| AATGTATGTGATTG | 39 |
| CCTGAACCGGATCCACTT | 39 |
| AGAAGA | 38 |
| CTGCTT | 38 |
| CTGCTA | 38 |
| CATTGC | 38 |
