## Extended Data Table 2 for "Human-like telomeres in *Zostera marina* reveal a mode of transition from the plant to the human telomeric sequences"

**Extended Data Table 2.** Predicted telomerase RNA sequences of Alismatales species. The sequences of the predicted telomerase RNAs were obtained from or ABG – Applied Bioinformatics Group (http://appliedbioinformatics.com.au/index.php/Seagrass_Zmu_Genome), NCBI - National Center for Biotechnology Information (https://www.ncbi.nlm.nih.gov/), and CoGe – Comparative Genomics (https://genomevolution.org/coge/).

TR1 and TR2 indicate two possible paralogs for telomerase RNA. TR1a and TR2 from *Z. marina* are genomic sequences. TR1a includes complete promoter and terminator non-transcribed sequences, while TR2 from *Z. marina* lacks several promoter elements. The TR1b from *Z. marina* was assembled for this study from reads obtained in RNA-Seq from root tips. Underlined nucleotides represent the predicted template regions. Note, the red dinucleotide of CT in *Z. marina* TR1b, which as an evolutionary novelty changes the template annealing potential to the preferred human-like telomere sequence. On the other hand, the potential for the plant telomere motif synthesis has not been completely lost in *Z. marina* template motif (green highlighting). In the non-transcribed TR2 of *Z. marina*, there is an additional mutation (C --> T) in the template region, which would potentially lead to (TTAGG)n (known from insects, blue highlighting) telomere sequence. From this point of view, the *Z. marina* TR represents transitional state producing human-type sequence, with a trace of plant-type synthesis.

| |
|---|
| ***Lemna minor* (TR1) (Lemnaceae)**<br>Sequence ID: lminor_contig_4989, Coordinates: 12166-12483, Source: CoGe, Genome ID:27408 |
| AATCCCACATCGGAAAATCAGAGAAATCAAGATTTAATATATTTGCAAGTCTGCTAAAAATAGTAATTATTCCAGGGGAAGGGTG<br>AGAGAGGTTAGAGAGAGAAACTGCTACTGAGTAAACCCTAAACCGTACTCTTAATTGAGGAATCTACCGGGCTTGATAGTGGGC<br>TGTTTGTCCGGCGTTTGAGCTCCCGGGTTGAAAGGCCCAATGAAATGCCGATGCACGCGGCTTTCTCTCTCTAAACCATTGGAAG<br>AGGCTTGTAGGGGGCAAACTGATTTGCCGATTTCCCTCGTCCCTCCCAAAATCCCCTGTTTTCTTTC |
| ***Lemna minor* (TR2) (Lemnaceae)**<br>Sequence ID: lminor_contig_4989, Coordinates: 10338-10654, Source: CoGe, Genome ID:27408 |
| AATCCCACATCGAAAAATCAGAGTAATTGAGATTCATTATATTTCGATGCCTGGTTAAAAAAACAATAATTCCAGGGGAACGATG<br>AGAGAGATTAGAGAGAGGAATTGCTACTGGTTAAACCCTAAACCGTACTCTTCATTGAGGATTCAGTTGGGCTTTGTTAACGGGC<br>TGTTTATCCGGCGTTTGAGCTCCCGGGTTGAAAGGCCCAATGAAACGCCGATGCACGCGGGTTTCTCTCCCTAATCCTTGGGAAG<br>AGGCTAGTAGGGGGCAAACTTGTTTGCCGATTTCCCTCGCCCTCCCAAATTCCCCTGTTTTCATAA |
| ***Posidonia oceanica* (TR1) (Posidoniaceae)**<br>Sequence ID: GFJT01022271.1 Coordinates: 1-297 Source: NCBI |
| TTCCATTTCCGCCCAAGCTGCTCACAGTATACGGGCGTCGGCATCCACCTTCCGGGGATGAGGTGGCTTGGGAGTACACTTGCAC<br>CAAAGCATGCTTATGTGTGCTCAAACCCTAAACTTTCCTCTTAAGTGAGGTTCGAGTTAACCTTACCAAAGAGGTACTTTGTCCAG<br>TATTTAACCTCTGTAATAAAAAAAATACTGATGCCTTGCACTTGTACTCCCAAAGTAGATATGCTACGGGAAGGCTTGAAGGGGG<br>TTGCCTCGGCAACCGATATCCTCGCCTTCCAAGTCCCCATT |

***Spirodela polyrhiza* (TR1) (Araceae)**

Sequence ID: UNPA01000014.1   Coordinates: 2999451-2999833  Source: NCBI

CCAAAATCCCACATCGGTAAATTTTTCAAGAAACATATGGGTTATATATTCAGCTGACCATTGGTTTTAATCTTCCAGGGGCCAAG
TGGGGATAGGAGGGCAGCCGCTCAGGCACCCGTTGACCGGGTTGTCCGATAAACCCTAAACCTGACTCTACTGAGGTCGACGCT
CCCAGCTTCACTTGGGTAGTGGAAAGAGGGCGGGGGCTGTTTACCCGGCGTTTAAGCCTCCCTTCTCCACCCGTCTTTGAGAAAT
TCAAACGCCGATGCCCGCGGCTGCCCCTCCCATCGATTATATCGTACCACGGGAGCGCTCGCAGGGGGCGGCGCCGCCAGGCTA
GCCGAGGTCCCTCGCCCTCCCAAGCCCCTGTTTTCTCTCTTCGA

***Zostera capensis* (TR1)**

Sequence ID: PRJNA503110 - assembled *de novo* from raw reads   Source: NCBI

GTCCCACACAGCCGTTGAGATGAAAAATAGAAGCATATATAAAACTAAAAAGCAAGACATACAACACCTCCTTGGGGGAATTTCA
CAAGGAGATTATTGCAGCAATCCCATCAGCAACATCGTTGGGATTAGAACCCTAAACCTACTTCTCTGGAAGGTCTTTAACGTCCA
CTGTACTGAGAAAGGACTGAGGAGGTACTTTTTCCGGTGTTTAACCTTCTATAAACAATAAAAACATCGATGCCCCTGCCTCTGTT
CTCCTGTGGGTTGTTGTTGGGTAAGTCTTGAAGGGGTTGTTTATGAAAAAAACACCGATGTCCTCGACTCTTCCCAACCCCAAGTT
CCCCTTTTTTATTTCAA

***Zostera marina* (TR1a)**

Sequence ID: LFYR01000762.1   Coordinates: 70571-71209   Source: NCBI (genome assembly by Olsen et al. 2015)

GTCCCACACCGCCAAAGAAATAAATTTTAAAGCATATATATAATAAAAAAACCTATTTATAAAATAACTTATTGGAGGGTTTGTGT
TTCCACCAGTAGTTTATTGCAGTTTGTCTATTCTAACCCTAAATACTTCTTTCTTCTAGAAAAGAGTTAATGTATGCTAAAAAGTAG
GTATTTTCTCCGAGATTTCTACCTGAAATGAATATGAAAAATCTCGATGCTTCTGCCTCTGTACTTCTGTGATTGATTGTTGGGGAA
ATCTTTGTAGGGAGTATTGTTTGTTCATAACTGATATCCTCGGCTTTCCTACAACAATCAAAATTCCCTCTGTTTTCATTTAATTT

***Zostera marina* (TR1b)**

Sequence ID: SRR10664318 – assembled *de novo* from raw transcriptomic reads Source: This study

GTTTGTGTTTCCACCAGTAGTTTATTGCAGTTTGTCTATTCTAACCCTAAATACTTCTTTCTTCTAGAAAAGAGTTAATGTATGCTAA
AAAGTAGGTATTTTCTCCGAGATTTCTACCTGAAATGAATATGAAAAATCTCGATGCTTCTGCCTCTGTACTTCTGTGATTGATTGT
TGGGGAAATCTTTGTAGGGAGTATTGTTTGTTCATAACTGATATCCTCGGCTTTCCTACAACAATCAAAATTCCCTCA

***Zostera marina* (TR2)**

Sequence ID: LFYR01001054.1   Coordinates: 856312 - 856652   Source: NCBI (genome assembly by Olsen et al. 2015)

CATGATATACATCCTTACCGGAGGGTTTGTGTATTGATCATAAGTACGTAGTTTATTACAGTTTGTTTCCACCAGTCGGTATAGGT
GGTATACTAACCTTAAATTCTCCTTTCTTTTAGAAAAGAGTTGATTTATGTAAAAATCAGATAGTTTCTTTGAGATTTATGTCTGAA
ATAAATATGAAAAATCTCGATGCTTCTGCCTCTGTAATCGATCTTGTCATTGATTCTTGGGGAAAGTTTAATTTGTAAGGAGTGAT
GCTTGTTCATCACTGATATATATCCTCGGCTTTCCTACAACAATCAAATTCCCTCTGTTTTTGATGTCAGAATATACGATTA

***Zostera noltii* (TR1)**

Sequence ID: SRR2409708 – assembled from raw reads Source: NCBI

CTTGGGGGAATTTACAAGGAGATTATTGCAGCAATCCCATCAGCAACATCGTTGGGATTAGAACCCTAAACCTACTTCTCTGGAA
GGTCTTTAATGTCCANNTGTATGAGAGAGGACTGAGGAGGTACTTTTTCCGGTGTTTAACCTTCTATAAACAATAAAAACATCGA
TGCCCCTGCCTCTGTTCTCCTGTGGGTTGTTG

***Zostera muelleri* (TR1)**

Sequence ID: Zmu_v1_s5374_62084   Coordinates: 11965-12335   Source: ABG

AGTCCCACACAGCCGTTGGGAAGAAAAATAGAAGCATATATAAAACAAATAAGCAAAATTTACATCACTTTCTTGGGGGGAATTT
CACTAGGAGACCAATGCAGCTTTCCCACCAGCAACATCGGTGGGATTAAACCCTAAACCAACTTCTTCGGAAGGTCTTTGATCAT
GTCCATTGTACTGGAGAGGACTGAGGAGGTACTTTTACCGGTGTTTAACCTTCTATAAACAATAAAAACATCGATGCCCCTGCGTC

TGTGTTCTCCAGTGGGTTGTTGGTTGGGTTAAGTCTTGAAGGGGGTGTTTATGAAAAAACACCATTGTCCTCGACTCTTCCCAACC
CCAAATTCCCCTCTTTTAATTTAATTTAA

***Zostera muelleri* (TR2)**

Sequence ID: Zmu_v1_s7592_29784_9366_29784   Coordinates: 8581-8221   Source: ABG

GTCCCACACAGCCGTTGGGATAAAAAATAGAAGCATATATAAAACTCAAAAGCAAGACTACGACACCTTCTTGGGGGAATTTCAC
AAGGAGATTATTGCAGCAATCCCATCAGCAACATCTTTGGGATTAGAACCCTAAACCTACTTCTTTGGAAGGTCTATAATGTCCAC
CGTACTGAGAGAGGACTGAGGAGGTACTTTTTCCGGTGTTTAACCTTCTATAAACAATAAAAACATCGATGCTCCTGCCTCTGTTC
TCCTGTGGGTTGTTGTTGGGTAAGTCTTGAAGGGGTTGTTTATGAAAAAACACCGATGTCCTCGACTCTTCCCAACCCCAAGTTCC
CCTCTTTTAATTTAATTT
